# Neuronal activity in sensory cortex predicts the specificity of learning

**DOI:** 10.1101/2020.06.02.128702

**Authors:** Katherine C. Wood, Christopher F. Angeloni, Karmi Oxman, Claudia Clopath, Maria N. Geffen

## Abstract

Learning to avoid dangerous signals while preserving normal responses to safe stimuli is essential for everyday behavior and survival. Fear learning has a high level of inter-subject variability. Following identical experiences, subjects exhibit fear specificities ranging from high (specializing fear to only the dangerous stimulus) to low (generalizing fear to safe stimuli). Pathological fear generalization underlies emotional disorders, such as post-traumatic stress disorder. The neuronal basis of fear specificity remains unknown. Here, we identified the neuronal code that underlies inter-subject variability in fear specificity using longitudinal imaging of neuronal activity before and after differential fear conditioning in the auditory cortex of mice. Neuronal activity prior to, but not after learning predicted the level of specificity following fear conditioning across subjects. Stimulus representation in auditory cortex was reorganized following conditioning. However, the reorganized neuronal activity did not relate to the specificity of learning. These results present a novel neuronal code that determines individual patterns in learning.

## Introduction

Learning allows our brain to adjust sensory representations based on environmental demands. Fear conditioning, in which a neutral stimulus is paired with an aversive stimulus, is a robust form of associative learning: exposure to just a few stimuli can lead to a fear response that lasts over the subject’s lifetime ^1,2^. However, the same fear conditioning paradigm elicits different levels of learning specificity across subjects ^3–6^. In pathological cases, the generalization of the fear response to stimuli in non-threatening situations can lead to conditions such as post-traumatic stress disorder (PTSD) ^7,8^ and anxiety ^9^. Therefore, determining the neuronal basis for learning specificity following fear conditioning is important and can lead to improved understanding of the neuropathology of these disorders. Whereas much is known about how fear is associated with the paired stimulus, the neuronal mechanisms that determine the level of specificity of fear learning remain poorly understood. Our first goal was to determine the neuronal basis for the differential fear learning specificity across subjects.

Multiple studies suggest the auditory cortex (AC) is involved in fear learning. *During* differential fear conditioning (DFC), inactivation of AC chemically ^10^, or with optogenetics ^11^, as well as partial suppression of inhibition in AC ^12^ led to decreased learning specificity using either pure tones or complex stimuli, such as FM sweeps or vocalizations ^3,11–14^. These observations suggest that AC may determine the level of learning specificity. Therefore, we tested whether neuronal codes in AC *prior* to conditioning can predict specificity of fear learning.

The role of AC *following* fear conditioning is more controversial. Changes in stimulus representation in AC following association learning have been proposed to represent multiple different features of the fear response ^1,14–19^. However, inactivation of the auditory cortex did not affect fear memory retrieval of pure tones ^3,11^, suggesting that AC is not involved in fear memory retrieval. If AC were involved in fear memory retrieval, we would expect the changes in sound representation to reflect the level of learning specificity across subjects. Therefore, our second goal was to test the role of changes in auditory cortex in shaping fear learning specificity across subjects.

To address these goals, we imaged the activity of neuronal ensembles in layers 2 and 3 of AC over weeks, before and after differential fear conditioning with pure tones. First, we established the neuronal basis for differential learning specificity across subjects by finding that neuronal activity in AC prior to fear conditioning predicted the level of learning specificity. Second, we found that the changes in stimulus representation in AC following fear conditioning were not correlated with the level of learning specificity across subjects, suggesting that the role of AC in fear learning is restricted to the consolidation period and changes in AC do not represent fear memory. These findings refine our understanding of the neuronal code for variability in fear learning across subjects and reconcile seemingly conflicting previous results on the function of the auditory cortex in fear learning.

## Results

### Learning specificity varies amongst conditioned mice

To establish the relationship between sound-evoked activity in the AC and differential fear conditioning, we recorded simultaneous neuronal activity from hundreds of neurons in AC. We tracked the same neurons before and after DFC, using two-photon imaging of a fluorescent calcium probe (GCaMP6^20^, Fig S1-2). Longitudinal imaging of neuronal activity in large ensembles of neurons in layers 2 and 3 of AC before and after conditioning (Fig 1a) allowed us to compare the representation of the CS stimuli before and after learning.

**Figure 1:**
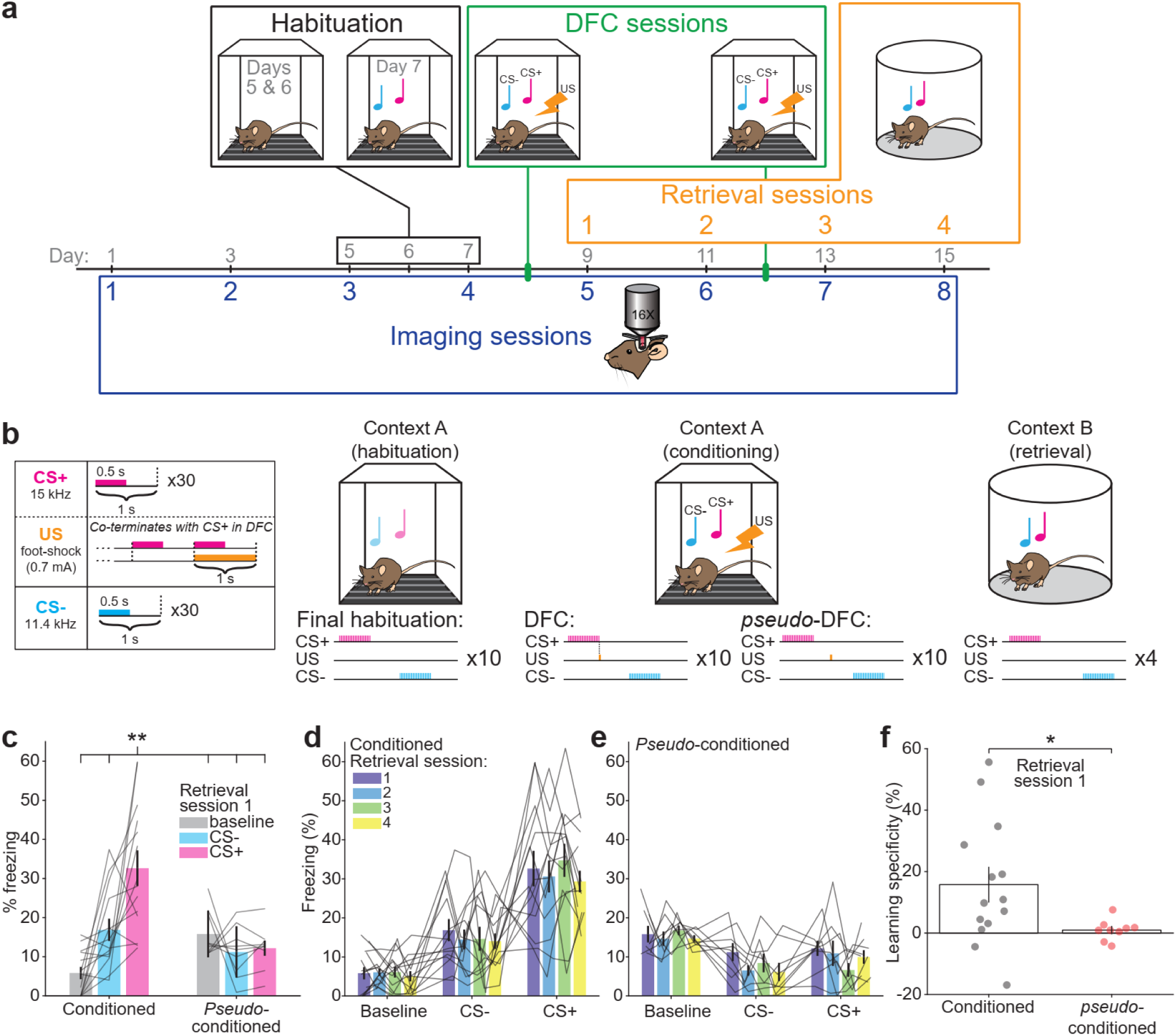
Experimental timeline and differential fear conditioning (DFC) paradigm. **(a)** Experimental timeline: Mice were imaged for 4 sessions (48 hours apart) before DFC to establish baseline responses to tone pip stimuli under the two-photon. Prior to DFC, mice were habituated to the fear conditioning chamber. Mice were subjected to DFC (19 mice) or *pseudo*-conditioning (9 mice) on Days 8 and 12. After DFC-1 (day 8), fear retrieval testing was performed after each imaging session. **(b)** Mice were habituated to the conditioning chamber (context A) for 3 days prior to conditioning and on the final day, the stimuli were presented without foot-shock. During conditioning, a foot-shock (1 s, 0.7 mA) was paired with the CS+ (15 kHz, 30s pulsed at 1 Hz). The CS- (11.4 kHz, 30 s pulsed at 1 Hz) was presented alternately with the CS+ (30-180 s apart, 10 repeats) and not paired with a foot-shock. During *pseudo-*conditioning, 10 foot-shocks were presented randomly between the CS stimuli. During retrieval testing (context B), the same CS+ and CS- stimuli were presented alternately (30-180 s apart, 4 repeats). Motion of the mouse was recorded and the percentage freezing during each stimulus was measured offline. **(c)** Freezing at baseline (gray), for CS+ (pink) and CS- (blue) in retrieval session 1 (day 9) showing the percentage of time frozen during tone presentation for CS+, CS- and baseline for each mouse. Gray lines indicate freezing for each included mouse. (Two-way ANOVA, *Tukey*-*Kramer post-hoc* test, Table S1). **(d)** Freezing to baseline, CS-, and CS+ for each conditioned mouse over the 4 retrieval sessions. Gray lines show each mouse. **(e)** Same as **d** for *pseudo*-conditioned mice. **(f)** Learning specificity of conditioned and *pseudo*-conditioned mice for retrieval session 1. Circles show individual mice. (*t*-test). Error bars in **c-f** indicate standard error of the mean (sem). ^†^*p* < 0.1, \**p* < .05, \*\**p* < .01, \*\*\**p* < 0.001, ^n.s.^*p* > .05.

We conditioned mice by exposure to 10 repeats of an alternating sequence of two tones, one of which co-terminated with a foot-shock (CS+, 15 kHz), and one which did not (CS-, 11.4kHz). *Pseudo*-conditioned mice were presented with the same stimuli, but the foot-shock occurred during periods of silence between the stimuli (Fig 1b). Following conditioning, we measured fear-memory retrieval by presenting the same auditory stimuli to the mice in a different context and measuring the percentage of time the mice froze during stimulus presentation and at baseline (Fig 1c). Memory retrieval was tested after each imaging session. To test whether levels of freezing changed over retrieval sessions we fit a linear mixed-effects model to predict how freezing was affected by the retrieval session time and stimulus type. We found there was no effect of retrieval session on freezing (Fig 1d, Table S1, *t*_(164)_ = 0.90, *p* = .372) and no difference in the effect between retrieval session and stimulus type (*t*_(164)_ = -1.21, *p* = .227). Similarly, freezing in *pseudo-*conditioned mice was consistent over the 4 retrieval sessions (Fig 1e, Table S1, no effect of retrieval session or retrieval session*stim_type, *p* > .05). Since there was no change in freezing over time, we do not specifically consider results with respect to the second DFC session (Fig 1a, day 12). Henceforth we refer to DFC as the first DFC session. Conditioned mice that did not freeze to CS+ or CS- differently from baseline were excluded from subsequent analysis (5/19 mice excluded, Fig S3a, two-way ANOVA, *p* > .05, see methods).

Learning specificity was defined as the difference between freezing to CS+ and CS- during memory retrieval sessions (see Methods, Equation 1)^3^. We used two pure-tone CS stimuli which have been shown to engage AC in both mouse^3,12^ and human DFC^21^. The pure tones were close together in frequency space (0.40 octaves apart) in order to drive a range of learning specificities in conditioned mice that are not achievable at greater frequency distances^3^. Indeed, we observed that conditioned mice displayed a larger range of learning specificities (range: -16.9 to 55.6%) compared with *pseudo*-conditioned mice (−4.2 to 7.6%). This was reflected in a significantly larger standard deviation of learning specificity in conditioned mice (*σ* = 20.3%) than in *pseudo*-conditioned mice (Fig 1f, *σ* = 3.3%, *F*-test, *F*_(13, 8)_ = 36.80, *p* < .001) in the first retrieval session after DFC. We also observed a significantly higher learning specificity (mean: 15.8%) in conditioned mice than *pseudo*-conditioned mice (mean: 1.0%, *t*-test, *t*_(21)_ = 2.15, *p* = .043) in the retrieval session after DFC. To test whether learning specificity was consistent over retrieval sessions, we fit a linear mixed-effects model to predict how learning specificity was affected by retrieval session and conditioning type. We found no effect of retrieval session on learning specificity for conditioned mice (Fig S3b-c, Table S1, *t*_(88)_ = 0.23, *p* = .817) nor any interaction between session and conditioning type (*t*_(88)_ = -0.01, *p* = .995). Thus, we found that conditioned mice exhibited a range of learning specificities, with some generalizing their fear across the CS stimuli and others specializing their fear responses to CS+. On average, the learning specificity of mice was stable over the course of the experiment.

### Neuronal responses in AC pre-DFC predict specificity of fear learning

We used two-photon imaging to record calcium activity from neurons in auditory cortex in head-fixed mice (Fig 2a). We presented 100-ms tone pips (frequency range: 5-32 kHz, including CS+ and CS- frequencies) to obtain frequency response functions from each neuron. We hypothesized that the activity in auditory cortex would predict learning specificity across individual mice. Thus, we tested whether neuronal discrimination of CS+ and CS- in AC pre-DFC predicted learning specificity following DFC. To assess how well single neurons could discriminate between the two conditioned tones, we computed the Z-score difference (Z_diff_) of responses to CS+ and CS- for responsive neurons (see Methods, Equation 2). In an example neuron (Fig 2b), the distributions of single-trial response magnitudes to CS+ and CS- demonstrate a separation resulting in a significant Z_diff_ score of 2.01. The Z_diff_ score of responsive neurons was considered significant if the actual score was greater than the 95^th^ percentile of the bootstrapped Z_diff_ scores (see Methods). Figure 2c shows the distribution of Z_diff_ scores for all responsive units from conditioned mice 24 hours pre-DFC.

**Figure 2:**
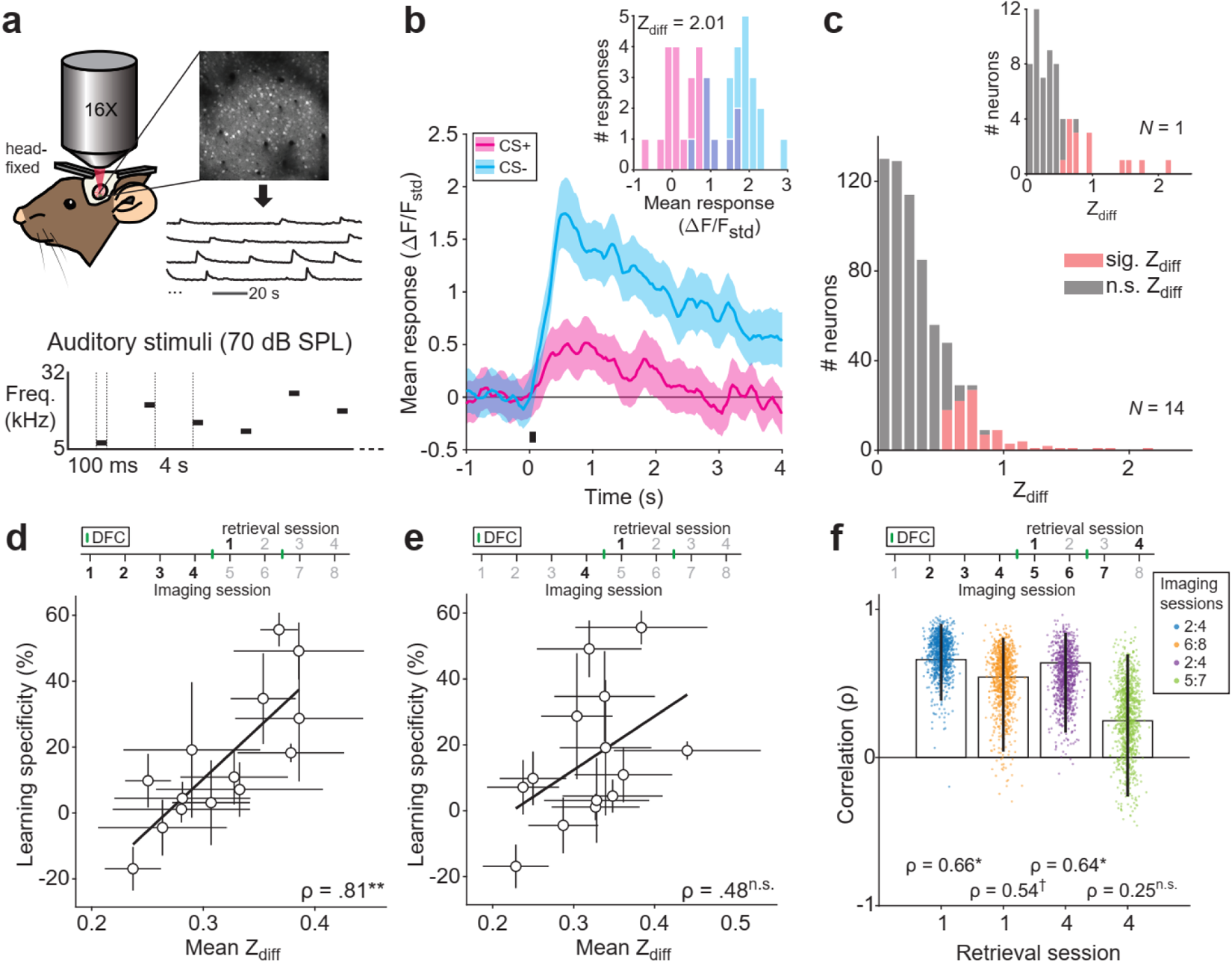
Mean neuronal discriminability pre-DFC predicts learning specificity. **(a)** Imaging setup: Mice were head-fixed under the two-photon microscope, fluorescence of calcium indicator (GCaMP6s/m) was measured at ∼30 Hz, regions of interest and mean fluorescence over time were extracted using open software^22^. Schematic showing auditory stimuli, comprised of pure-tone pips (100 ms, 70 dB SPL, 5-32 kHz) presented at 0.24 Hz. **(b)** Response (mean ± sem, 25 repeats) to the presentation (black bar) of CS+ (magenta) and CS- (cyan) of an example neuron. Inset shows distributions of the single-trial mean responses (mean ΔF/F_std_ across 2-s window following stimulus onset) to CS+ and CS- from the same neuron. **(c)** Distribution of Z_diff_ scores of responsive units from conditioned mice 24 hours pre-DFC. Significant scores are indicated in red, *n* = 98/653 neurons. Inset, single mouse example, *n* = 15/63 neurons. **(d)** Mean Z_diff_ (± sem) pre-DFC correlated (Spearman’s rank correlation, *N* = 14 mice) with learning specificity 24 hours post-DFC. Black line = linear best fit. **(e)** Mean Z_diff_ (± standard deviation [sd]) for each mouse 24 hours pre-DFC did not correlate with learning specificity 24 hours post-DFC. black line = linear best fit, *N* = 14 mice. **(f)** Spearman’s rank correlation (*ρ* ± 95% CI, *N* = 14 mice) between Z_diff_ score averaged across 3 imaging sessions as indicated in the legend and learning specificity from retrieval sessions 1 and 4. Dots represent individual bootstrapped *ρ* (*n* = 1000). ^†^*p* < 0.1, \**p* < 0.05, \*\**p* < 0.01, \*\*\**p* < 0.001, ^n.s.^*p* > 0.10.

To test whether neuronal discrimination pre-DFC could predict subsequent learning specificity, we averaged the Z_diff_ scores of neurons recorded in each recording session of each mouse and compared it with learning specificity 24 hours post-DFC. Since different numbers of neurons were recorded from each mouse, we resampled (100x with replacement) the lowest number of neurons recorded from across the mice. We found that the mean Z_diff_ scores averaged across the pre-DFC imaging sessions predicted learning specificity 24 hours post-DFC (Fig 2d, Spearman’s rank correlation, *ρ*(12) = .81, 95% confidence intervals (CI) [.60, .93], *p* = .007). Using only the imaging session preceding DFC, the mean Z_diff_ did not correlate with learning specificity 24 hours post-DFC (Fig 2e, *ρ*(12) = .48, 95% CI [.04, .78], *p* = .103). However, the two correlations were not significantly different from one another (bootstrap comparison, see methods: *ρ* difference = 0.33, 95% CI [-0.08, 0.88], *p* = 0.128, *N* = 14). In summary, this suggests that the neuronal discriminability in AC of individual mice pre-DFC predicts learning specificity 24 hours post-DFC.

It is possible that the Z_diff_ score results from some underlying distributions of response magnitudes; for example, the magnitude of response to CS+ could be driving the prediction phenomenon. Thus, we explored whether magnitude of CS+ or CS- responses related to learning specificity. We compared the mean response magnitudes to each CS over the 4 pre-DFC imaging sessions with learning specificity 24 hours post-DFC and found that they were not correlated (Spearman’s correlation, *p* < 0.05, Fig S4). This suggests that it is not merely the magnitude of responses to CS+ or CS- but truly discriminability of the responses that is underlying the prediction of learning specificity.

We next tested the temporal window for the prediction of learning specificity. If changes in sound-evoked responses in AC following DFC reflect memory formation or the strength of learning, as previously suggested^15,17^, we would expect a stronger relationship between neuronal discrimination and learning specificity after DFC than before. To test this, we compared the correlations between mean Z_diff_ across equal numbers of imaging sessions before and after DFC (3 imaging sessions preceding retrieval sessions 1, and 4) and learning specificity in retrieval sessions 1 and 4, respectively. We found that the mean Z_diff_ score pre-DFC predicted learning specificity from retrieval session 1 (Fig 2f blue dots, *ρ*(12) = .66, 95% CI [.39, .90], *p* = .031), whereas the mean Z_diff_ score post-DFC did not predict learning specificity in retrieval session 4 (Fig 2f green dots, *ρ*(12) = .25, 95% CI [-.26 .70], *p* = .401). However, these two correlations were not significantly different (bootstrap comparison (see Methods): *ρ* difference = 0.41, 95% CI [-0.09, 0.95], *p* = .124). This change in prediction could result from a rearrangement of learning specificity over time or a rearrangement of Z_diff_ scores over time. We reasoned that if learning specificity was rearranged then the neural discriminability pre-DFC ought not to correlate with the learning specificity in retrieval session 4 (Fig 2f purple dots). However, we found that neural discriminability pre-DFC was able to predict learning specificity in retrieval session 4 (*ρ*(12) = .64, 95% CI [-.17 .84], *p* = .022), suggesting a rearrangement of Z_diff_ scores. Further supporting a rearrangement of Z_diff_ scores, neuronal discriminability post-DFC did not correlate with learning specificity in retrieval session 1 (Fig 2f orange dots, *ρ*(12) = .54, 95% CI [-.04 .81], *p* = .066).

To verify that the results were robust to variability in frequency tuning distributions and location of the imaging window along the anterior-posterior axis (Fig S1 & S2) between mice, we investigated the relationship between Z_diff_ and these parameters. If neuronal discriminability is affected by imaging location then we would expect a relationship between the location of the imaging field of view on the anterior-posterior axis and Z_diff_, we did not find any relationship between these two measures (Fig S5a, Spearman’s rank correlation, *p* > .05) nor between the percentage of neurons with significant Z_diff_ and imaging location (Fig S5b). The best frequency distributions of neurons in the imaging window could affect the mean Z_diff_ of neurons of each mouse, if so we would expect higher Z_diff_ scores and more neurons with significant Z_diff_ scores for neurons tuned around the CS+ and CS-. However, we found no relationship between mean Z_diff_ score and mean best frequency in the imaging window (Fig S5c, Spearman’s rank correlation, *p* > .05) nor between the percentage of significant Z_diff_ scores and mean best frequency (Fig S5d). Not surprisingly, the percentage of significant Z_diff_ scores was correleated with learning specificity (Spearman’s rank correlation, *ρ* = .72, *p* = .011) suggesting that the best discriminating mice also had more neurons that discriminated between CS+ and CS- (Fig S5e). While there was no relationship between mean Z_diff_ and mean best frequency of all neurons in the imaging window across mice, we did find that neurons with best frequency at CS+ or CS- had higher Z_diff_ scores than neurons tuned to other frequencies (Fig S5f, Table S1). This suggests that mice with more neurons with best frequencies at CS+ and CS- might have better learning specificity. However, there was no relationship between the percentage of neurons in the imaging window with best frequency at CS+ and CS- across the pre-DFC imaging sessions and learning specificity (Spearman’s rank correlation, *ρ*(12) = .46, 95% CI [-.06, .71], *p* = .127).

In summary, individual neuronal discriminability in AC pre-DFC predicted learning specificity 24 hours after DFC. Post-DFC, neuronal activity no longer predicted learning specificity. Therefore, the role of auditory cortex in DFC is likely restricted temporally. To further investigate the relationship between neuronal and behavioral discriminability, we examined whether neuronal population discriminability could predict learning specificity.

### Population neuronal activity in AC predicts specificity of learning

For many brain regions and tasks, activity of multiple neurons can provide more information in combination than averaged activity of individual neurons^23–25^. Using machine learning, we investigated whether populations of neurons predicted learning specificity better than the average Z_diff_ scores. We trained a Support Vector Machine (SVM) to discriminate between presentation of CS+ and CS- using population responses to the two stimuli – again we resampled (100x with replacement) the lowest number of neurons recorded from across the mice. Mean SVM performance across imaging sessions prior to DFC correlated with learning specificity 24 hours post-DFC (Fig 3a, *ρ*(12) = .77, 95% CI [.53, .89], *p* = .001). Using only the imaging session preceding DFC, SVM performance 24 hours pre-DFC did not correlate with learning specificity 24 hours post-DFC (Fig 3b, *ρ*(12) = .35, 95% CI [-.20, .64], *p* = .247). However, the two correlations were not significantly different (bootstrap comparison (see Methods), *ρ* difference = -0.43, 95% CI [-0. 86, 0.00], *p* = .056). The Z_diff_ scores and the SVM performance of the same neurons were strongly correlated (Fig 3c, *ρ*(12) = .93, 95% CI [.89, .97], *p* < 0.001), suggesting that the two different discriminability methods used similar underlying features to discriminate the stimuli. This was also reflected in the fact that the correlations between the two discriminability measures across pre-DFC imaging sessions and learning specificity were not statistically different (bootstrap comparison*, ρ* difference = 0.01, 95% CI [-0.09, 0.00], *p* = .780). Thus, population responses averaged across pre-DFC imaging sessions predicted subsequent learning specificity likely through similar mechanisms to the mean Z_diff_. Since the SVM should give greater weight to more informative neurons, we tested whether there would be a stronger correlation between the *significant* Z_diff_ scores and SVM performance. We found that the correlations were not significantly different (bootstrap comparison*, ρ* difference = -0.044, 95% CI [-0.28, 0.14], *p* = .562).

**Figure 3:**
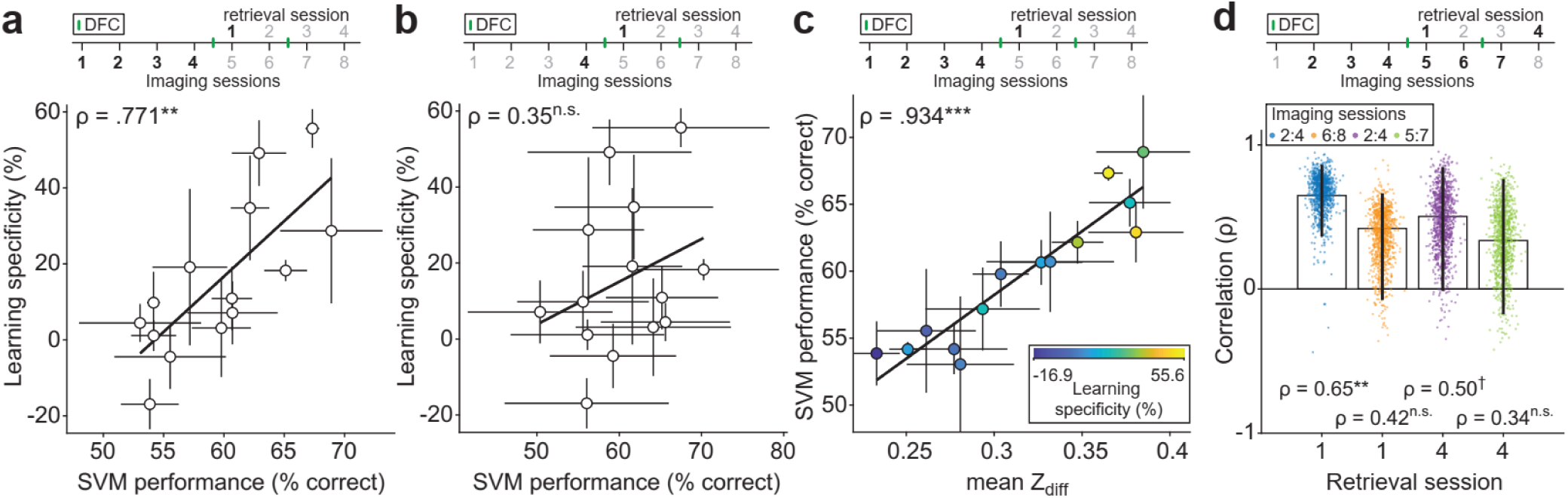
Neuronal population discrimination between CS+ and CS- pre-DFC predicts learning specificity. **(a)** SVM performance (mean ± sem) across pre-DFC sessions predicts learning specificity 24 hours post-DFC (retrieval session 1). **(b)** SVM performance (mean ± sd) 24 hours pre-DFC does not predict learning specificity 24 hours post-DFC. **(c)** Mean (± sem) SVM performance pre-DFC correlates with the mean (± sem) Z_diff_ score pre-DFC. Fill color indicates learning specificity from retrieval session 1. **(d)** Correlation (*ρ* ± 95% CI) between SVM performance averaged across 3 imaging sessions preceding retrieval sessions 1 (blue), and 4 (orange). Dots represent individual bootstrapped correlation values (*n* = 1000). Black lines in **a**, **b**, & **c** show the best linear fit. Statistics in **a-d**: Spearman’s rank correlation. ^†^*p* < 0.1, \**p* < 0.05, \*\**p* < 0.01, \*\*\**p* < 0.001, ^n.s.^*p* > 0.10.

We next tested whether predictability of learning specificity persisted after DFC by comparing the mean SVM performance across groups of 3 imaging sessions with retrieval sessions 1 and 4 (Fig 3d). The mean SVM performance pre-DFC predicted learning specificity in retrieval session 1 (Fig 3d blue dots, *ρ*(12) = .65, 95% CI [.36, .86], *p* = .008) whereas the mean SVM performance post-DFC did not predict learning specificity in retrieval session 4 (Fig 3d green dots, *ρ*(12) = .34, 95% CI [-.18, .76], *p* = .252). However, these two correlations were not significantly different (bootstrap comparison, *ρ* difference = 0.31, 95% CI [-0.24, 0.93], *p* = .294). Again, we tested whether this change in prediction resulted from a rearrangement of learning specificity over time or a rearrangement of neural discrimination over time. While we found the same pattern of results as in the Z_diff_ (Fig 2f), we did not find a significant correlation between neural discriminability pre-DFC and learning specificity in retrieval session 4 (Fig 3d purple dots, *ρ*(12) = .50, 95% CI [-.01 .85], *p* = .073). Neuronal discriminability post-DFC did not correlate with learning specificity in retrieval session 1 (Fig 3d orange dots, *ρ*(12) = .42, 95% CI [-.08 .66], *p* = .155).

Combined with similar results from the mean Z_diff_ scores (Fig 2f), there is evidence to support that neuronal discriminability predicts learning specificity *before, but not after,* conditioning. This is consistent with the hypothesis that neural activity is reorganized following DFC and that auditory cortex can no longer modulate the freezing response following conditioning, as suggested by previous work showing that learning specificity is not dependent on auditory cortical activity after fear conditioning (Aizenberg & Geffen 2013). Since neuronal activity no longer predicts learning specificity after conditioning, we hypothesized that there would be changes in neuronal activity following conditioning. Therefore, we next investigated changes in response and neural discriminability following DFC.

### After DFC, neuronal discriminability between CS+ and CS- is preserved

It has been suggested that ‘fear memories’ are encoded in the auditory cortex following differential fear conditioning^15,17^, implying that neuronal discriminability may improve following conditioning. We found that neuronal activity following DFC no longer predicted learning specificity (Fig 2f & 3d), suggesting AC does not support the fear response after DFC. We tested whether the neuronal discriminability of CS+ and CS- changed after DFC by comparing the mean Z_diff_ across pre- and post-DFC sessions (Fig 4a). We found no change in Z_diff_ from pre- to post-DFC in conditioned mice (Table S1, rm-ANOVA *Tukey-Kramer post-hoc* comparison, *p* = .740), whereas there was a significant decrease in *pseudo-*conditioned mice (*Tukey-Kramer post-hoc* comparison, *p* = .028). Results were similar at a neuronal population level; mean SVM performance in conditioned mice did not change across pre- and post-DFC sessions (Fig 4b, Table S1, rm-ANOVA *Tukey-Kramer post-hoc* comparison, *p* = .573), whereas there was a significant decrease in *pseudo*-conditioned mice (*Tukey-Kramer post-hoc* comparison, *p* = .001). Combined, we found that following DFC or *pseudo*-conditioning, neuronal discrimination between the CS+ and CS- was maintained in conditioned mice, while it decreased in *pseudo*-conditioned mice. These results suggest that changes in AC do not improve neural discriminability . Rather, plasticity in AC in conditioned mice appeared to counteract previously reported habituation in neuronal responses to repeated stimuli^18,26^.

**Figure 4:**
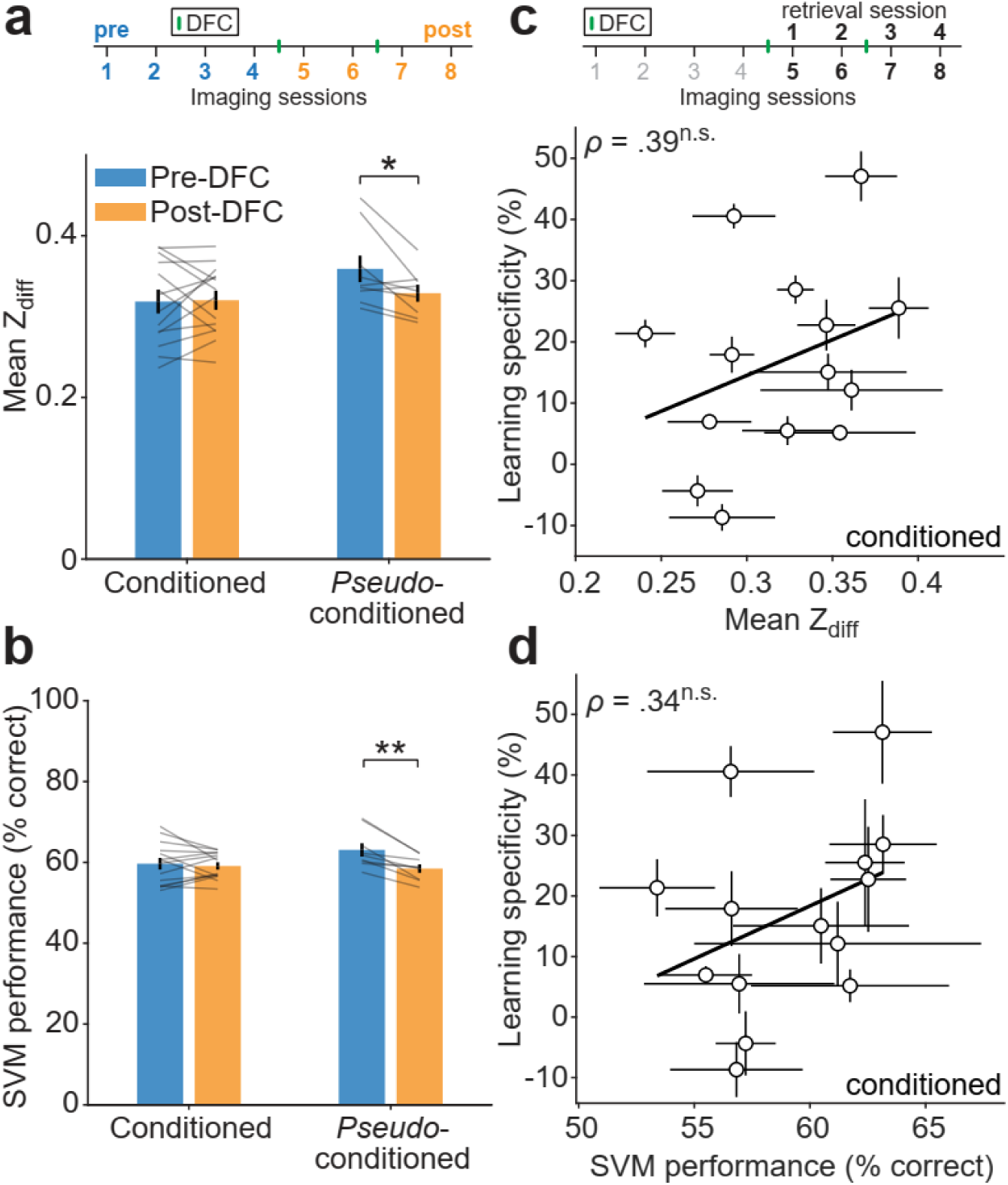
Changes in neuronal discrimination post-DFC. **(a)** Comparison of mean ± sem Z_diff_ between the pre- (sessions 1-4, blue) and post-DFC sessions (5-8, orange) in conditioned and *pseudo*-conditioned mice. Statistics: *Tukey-Kramer post-hoc*, Table S1. **(b)** Same as **a** but for comparison of mean ± sem SVM performance between the pre- and post-DFC. Stats: *Tukey-Kramer post-hoc*, Table S1. **(c)** Relationship between mean Z_diff_ across the post-DFC sessions (sessions 5-8) and mean learning specificity across all retrieval sessions. Statistics: Spearman’s rank correlation. Black line shows best linear fit. **(d)** Same as **c** but for mean SVM performance across the post-DFC sessions and mean learning specificity. Statistics: Spearman’s rank correlation. ^†^*p* < 0.1, \**p* < 0.05, \*\**p* < 0.01, \*\*\**p* < 0.001, ^n.s.^*p* > 0.10.

To further investigate how neuronal discrimination changed over time, we tested the neuronal discrimination performance of the SVM using cells tracked across pairs of imaging sessions. We trained the SVM using one imaging session and tested on data held out from that session and from the same cells in the second testing session (Fig S6a & b). If neuronal discriminability is maintained in conditioned mice, we would expect that there would be no change in performance between training and testing sessions. By contrast, in *pseudo*-conditioned mice, as neuronal discriminability appears to decrease, we expected to observe a decrease in performance particularly between sessions pre- and post-DFC. In conditioned mice, there was a small deficit in the testing sessions compared with training sessions, which did not change over sessions. By contrast, in *pseudo*-conditioned mice, we observed the same deficit in testing sessions compared with training, but the deficit increased over sessions. A linear regression of difference in performance with mouse group (m) and # sessions between testing and training (s) as predictors indicated that the slope of the relationship was significantly different between conditioned and *pseudo*-conditioned mice (Table S1, m*s, *p* = .020). Similarly, we observed a decrease in Z_diff_ as the number of sessions between pairs increased in *pseudo*-conditioned mice, but not in conditioned mice (Fig S6c & d, Table S1, Linear regression, m*s, *p = .*004). Neuronal representations are stabilized over time with behavioral relevance and drift without^26,27^. To assess whether representation of the CS+ and CS- was stabilized in conditioned vs. *pseudo*-conditioned mice we investigated whether was drift in the Z_diff_ of populations of neurons. If there is drift in the neuronal representation, then the similarity of Z_diff_ between individual neurons over time should become progressively dissimilar. We calculated the similarity (Pearson’s correlation) of Z_diff_ scores of neurons tracked between pairs of imaging sessions (Fig S6e). We fit a linear mixed-effects model to predict how Z_diff_ similarity between sessions was affected by the time between imaging sessions and whether mice were conditioned or *pseudo*-conditioned. We found there was a negative effect of number of imaging sessions between pairs of sessions on Z_diff_ similarity (Table S1, *t*_(624)_ = -2.87, *p* = .004) but no difference in the effect between conditioned and pseudo-conditioned mice (*t*_(624)_ = 1.28, *p* = .201). In summary, there is evidence of drift in the Z_diff_ score of both groups of mice, indicating that the Z_diff_ of individual cells became progressively dissimilar. In conditioned mice, the average Z_diff_ was maintained, while in *pseudo*-conditioned mice it decreased.

Different levels of learning specificity across mice could potentially account for the different levels of neuronal discriminability post-DFC. We therefore tested whether there was any correlation between the neuronal discriminability (mean Z_diff_ score and SVM performance) and the learning specificity *post-DFC*. The mean Z_diff_ score (imaging sessions 5-8) did not correlate with the mean learning specificity across retrieval sessions 1-4 of conditioned mice (Fig 4c, Spearman’s rank correlation, *ρ*(12) *=* .39, CI [-.25, .73], *p* = .175), nor was there a correlation between the mean SVM performance post-DFC and the mean learning specificity post-DFC (Fig 4d, *ρ*(12) = .34, CI [-.22, .79], *p* = .264). This suggests that neuronal discriminability post-DFC does not reflect learning specificity.

### After DFC, normalized responses at CS+ increased in conditioned mice

It has previously been shown that after differential conditioning with pure tones, select neurons in AC amplified the difference between CS+ and CS-^17,28^. However, since we observed no change in neuronal discrimination in conditioned mice, we hypothesized that there would be no change in response to CS+ and CS-. To test whether responses were altered by conditioning, we compared frequency response functions from the pre- and post-DFC imaging sessions of responsive neurons that were tracked from pre- to post-DFC (Fig S7a). On an individual neuron basis, we observed heterogeneous changes in the frequency tuning (Fig 5a, Table S1). However, on average, in conditioned mice, the *normalized* response to CS+ and frequencies between the CS+ and CS- increased, whereas the response at CS- did not change (Fig 5b, two-way rm-ANOVA, *Tukey-Kramer post-hoc* testing, *p* < .05, Table S1). In contrast, in *pseudo*-conditioned mice, the mean normalized responses at most frequencies, including both CS frequencies, did not change (Fig 5c, Table S1). When comparing normalized responses at CS- and CS+ in conditioned mice and the CS stimuli combined (CSc) in *pseudo-*conditioned mice, there was a significant increase at the CS+ and no change at CS- or CSc (Table S2, *Tukey-Kramer post-hoc* comparison, *p* < .001). Although we observed an increase in normalized response to CS+, there were no significant changes in non-normalized response to conditioned frequencies in conditioned mice (Fig S7b, d, & e, Table S1) and we observed decreased responses to most frequencies in *pseudo*-conditioned mice (Fig S7c, f, & g, Table S1). When comparing non-normalized response changes to CS+, CS- and CSc, we found a significant decrease at CSc but not at CS+ or CS- (Table S3, *Tukey-Kramer post-hoc* comparison, *p* < .001). It is possible that the normalization of the frequency response functions has amplified a small change that is not strong enough to present in the absolute responses.

**Figure 5:**
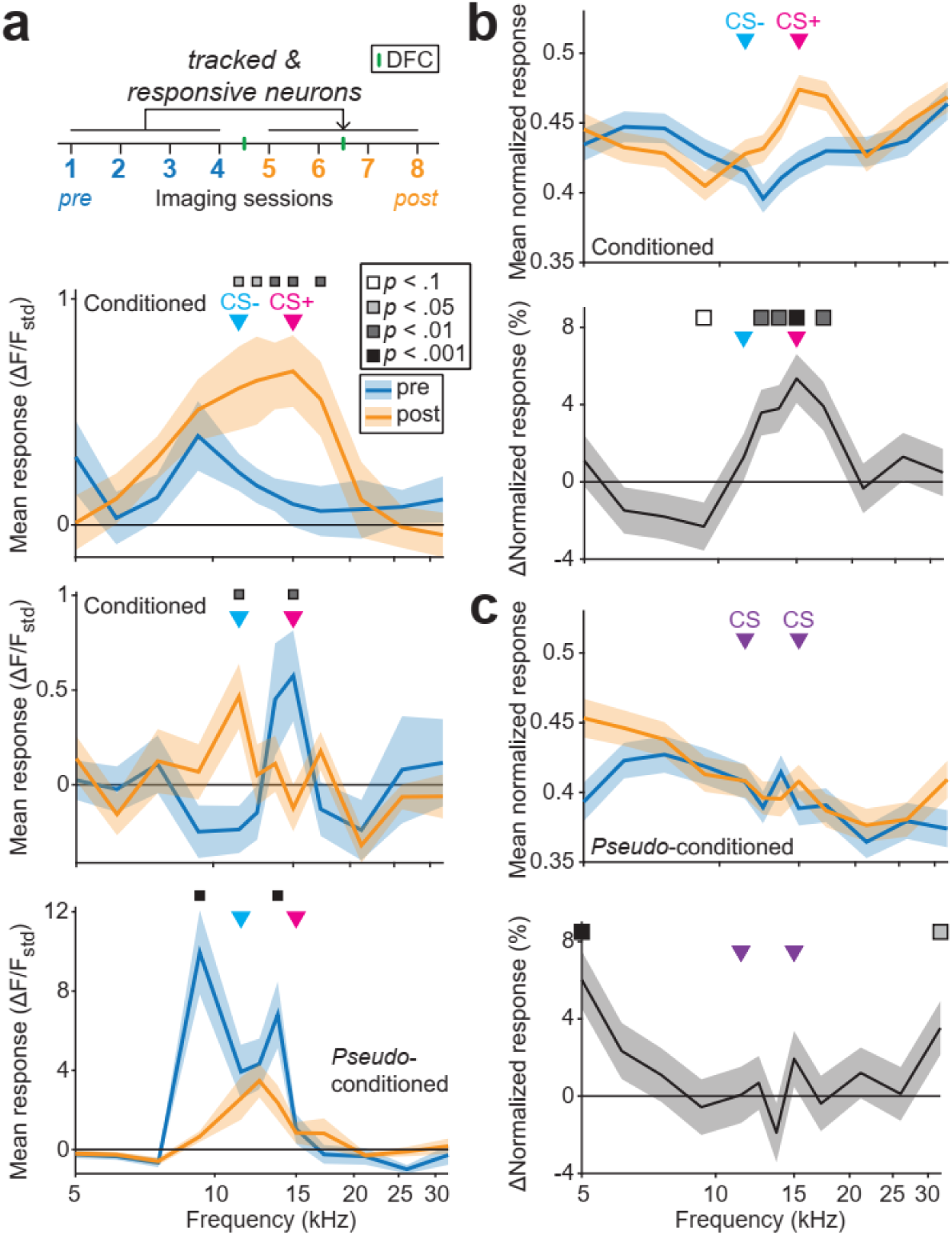
Changes in frequency representation post-DFC. **(a)** We tracked the responses of neurons responsive at least once pre- and post-DFC. The panels show three example frequency response functions from tracked neurons from conditioned and *pseudo*-conditioned mice pre-DFC (blue) and post-DFC (orange). Significant differences in the response functions are indicated by the squares above (two-way rm-ANOVA, *Tukey-Kramer post-hoc* analysis, Table S1). Arrows show the frequencies of the CS- (11.4 kHz) and CS+ (15 kHz) and CSc. **(b)** (top panel) Mean normalized frequency response functions of tracked responsive neurons across all conditioned mice (*N* = 14 mice, *n* = 879 neurons). (lower panel) Percent change in normalized frequency response functions of the same neurons, squares indicate significant changes (two-way rm-ANOVA, *Tukey-Kramer post-hoc* analysis, Table S1). **(c)** (top panel) Same **b** for pseudo-conditoned mice (*N* = 9 mice, *n* = 626 neurons). (lower panel) Percent change in normalized frequency response functions for the same cells as above, squares indicate significant changes (Two-way rm-ANOVA, *Tukey-Kramer post-hoc* analysis, Table S1).

Despite the lack of significant change in the absolute responses, it is possible that the increase in normalized responses at CS+ and the lack of change in response at CS- in conditioned mice could lead to improved discriminability between CS+ and CS- by increasing the difference between the responses to each stimulus. This would be consistent with the hypothesis that, following fear conditioning, reorganization of neuronal activity serves to amplify the *relative* difference in responses to CS+ and CS- thereby supporting discriminability^17,19^. However, when we compared the absolute normalized response post-DFC and the magnitude of changes in normalized response to CS+, CS-, and the difference between the two with learning specificity, we did not find any correlation (Fig S8), suggesting that the changes observed are in fact not related to storage of the fear memory^29^. We further investigated by checking for a relationship between the *change* in neuronal discrimination (Z_diff_ and SVM performance) and learning specificity (Fig S9) finding negative correlations between the two factors. This suggests that the neurons that were most predictive of learning specificity changed less than neurons that were less predictive, supporting the idea that reorganization of cortical activity following DFC does not depend on the fear memory, and may be due simply to random drift^27^.

Previous studies found that the best frequency of neurons shifts towards the conditioned stimulus (CS+) after DFC with pure tones^17^. We observed changes in the distributions of best frequencies following DFC (Fig 6a & b). To quantify the relationship of these changes to DFC, we calculated the absolute distance of the best frequency of responsive neurons to the CS+ frequency. Consistent with a shift in best frequency towards the CS+, we observed a small decrease in the absolute distance of best frequency from CS+ (of mean response functions pre- and post-DFC for each neuron) in responsive neurons of conditioned mice (Fig 6c, -0.07 octaves, two-way rm ANOVA, *Tukey Kramer post-hoc*, *p* < .001, Table S1) but not in *pseudo*-conditioned mice (*p* = .934). It is possible that neuronal discrimination between CS+ and CS- could be altered by a change frequency tuning width^12^. As a measure of tuning width we used the sparseness of the frequency response function^30,31^: A neuron with high sparseness responds strongly to one or few frequencies tested and little to other frequencies. A neuron with a sparseness of zero would indicate an equal response to all frequencies tested. We found no difference between the changes in the conditoned and *pseudo-*conditioned mice (Fig 6d, two-way rm-ANOVA, *F*_(1,1503)_ = 0.21, *p* = .649, Table S1) and that sparseness decreased in both (*F*_(1,1503)_ = 20.93, *p* < .001).

**Figure 6:**
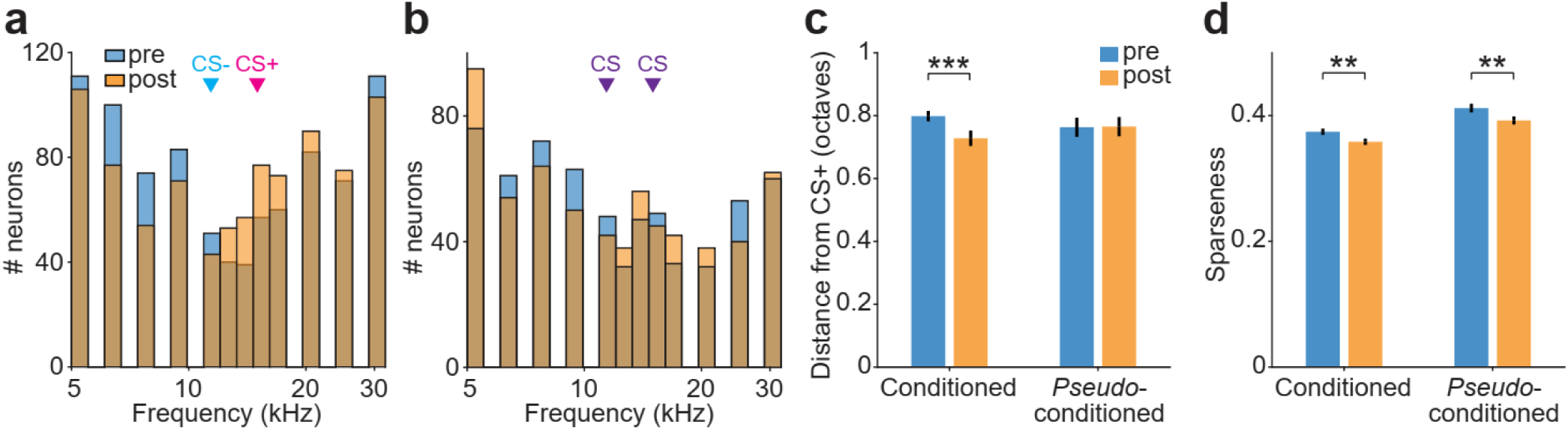
Best frequency and tuning sparseness pre- and post-conditioning. **(a)** Distributions of best frequencies of responsive neurons pre- (blue) and post-conditioning (orange), *N* = 879. **(b)** Same as **b** for *pseudo*-conditioned mice, *N* = 626. **(c)** Distance of best frequency from CS+ (15 kHz) of neurons from conditioned and *pseudo*-conditioned mice pre- (blue) and post-conditioning (orange). Statistics: 2-way rmANOVA *Tukey-Kramer post-hoc* analysis (Table S1). **(d)** Sparseness of mean frequency response functions of neurons from conditioned and pseudo-conditioned mice pre- (blue) and post-conditioning (orange). Statistics: 2-way rmANOVA *Tukey-Kramer post-hoc* analysis (Table S1). Error bars = sem. ^†^*p* < 0.1, \**p* < 0.05, \*\**p* < 0.01, \*\*\**p* < 0.001, ^n.s.^*p* > 0.10.

To verify that the results were robust to variability in frequency tuning between conditioned and *pseudo*-conditioned mice, we performed the analysis on change in response, change in distance of best frequency from CS+, and change in sparseness resampling the same number of neurons from each best frequency bin (1/12 bins). This had the effect of normalizing the frequency distributions pre-DFC between conditioned and *pseudo*-conditioned mice. We found that there was still an increase in response at the CS+ in conditioned mice while there were no changes at CS-, and no changes at either CS in *pseudo*-conditioned mice (Fig S10a). Furthermore, we found that despite the increase in response at CS+ in conditioned mice, there was no change in the distance of best frequency from the CS+ on average, whilst there was an increase in distance from CS+ in *pseudo*-conditioned mice (Fig S10b) while sparseness fell in both groups of mice (Fig S10c). Observing the best frequency distributions post-DFC (Fig S10d), this change is driven mostly an increase in neurons with best frequency at the extremes of our measurement (5 and 32 kHz). Qualitatively, the conditioned mice showed increased numbers of neurons tuned at and above the CS+ and decreased numbers below CS+ compared with *pseudo*-conditioned mice. Pairing of the CS+ with the shock led to increased number of neurons tuned to frequencies at and above the CS+ compared with an unpaired shock. Combined, whereas we find some changes in tuning consistent with classical results, these changes do not account for the differential learning specificity across mice.

To investigate whether variability in the region of sampling in each mouse affected the main findings, we split the mice into two groups based on whether the location that the center of their imaging window mapped onto the anterior-posterior axis. Locations that also contained the auditory thalamus (medial geniculate body, Fig S1) or not, with each group’s field of view more likely to be from primary auditory cortex (A1) or the anterior auditory field (AAF), respectively^32^. We found that the changes in response at CS+ were driven by neurons in putative A1 where there was a significant increase in normalized response and not in putative AAF where there was no change in response (Fig S11a). The distance of best frequency increased on average in AAF while there was no change in A1 (Fig S11b). However, we found no effect of imaging region on predicition of learning specificity by Z_diff_ or the SVM performance pre-DFC (Fig S11c-d) and no effect of imaging region on change in Z_diff_ (Fig S11e). Thus, there appears to be a differential effect of change in response at CS+ following conditioning for primary auditory cortex regions A1 and AAF, but this does not appear strongly related to the learning specificity (Fig S11f).

In summary, we observed heterogeneous changes in responses of individual neurons tracked from pre- to post-DFC. In conditioned animals, there was, on average, an increase in normalized response at CS+ and no change at CS-, however increase was not observed in absolute response changes. In *pseudo*-conditioned mice, we observed no changes in normalized responses at the CS stimuli. We observed a small shift in best frequency towards CS+ in conditioned mice. Sparseness of the frequency response functions decreased in both conditioned and *pseudo*-conditioned mice, indicating that frequency tuning became broader after conditioning, thus unlikely to support increased discriminability. Combined, these results reconcile our findings with previous studies, which had effectively, by not sampling responses from the same neurons pre- and post-DFC, normalized the responses. It is plausible that previous studies observed an increase in normalized activity, which did not translate into an actual population-wide increase in discriminability as we find here.

### A learning model of the fear circuit

We found that AC activity prior to learning predicts specificity of learning, yet the reorganized neuronal responses do not correlate with learning specificity. In order to better understand our findings in relation with previous results, we built a simple model that consisted of two frequency-tuned populations of neurons and a neuronal population that responds to the foot-shock. Our goal was to test whether this simple model could account for both the findings in this manuscript and from previous work, in particular: (1) Discriminability between CS+ and CS- in AC predicts learning specificity post-DFC (Fig 2-3); (2) Suppressing inhibition in AC leads to increased generalization (decreased learning specificity) post-DFC ^12^; (3) Suppressing AC post-DFC does not affect learning specificity ^3,11^.

In the model, we included two populations of frequency-tuned neurons (representing the medial geniculate body, MGB, and AC). MGB receives auditory inputs and projects to AC. Both populations project to basolateral amygdala (BLA). AC sends tonotopically organized feedback connections to MGB. During conditioning, the MGB neurons receive sound inputs and the neurons in the BLA are active during the foot-shock (Fig 7a). The weights from MGB and AC to BLA are updated according to a Delta learning rule (see Methods), that is, they are potentiated when both are co-activated (i.e. when the foot-shock coincides with the sound stimulus). We control the level of overlap in frequency tuning between neurons in AC, which difts over time^27^, and use it to represent frequency discriminability (more overlap = less discriminability). The activity of the BLA after weight update and with auditory input only is used as a measure of freezing.

**Figure 7:**
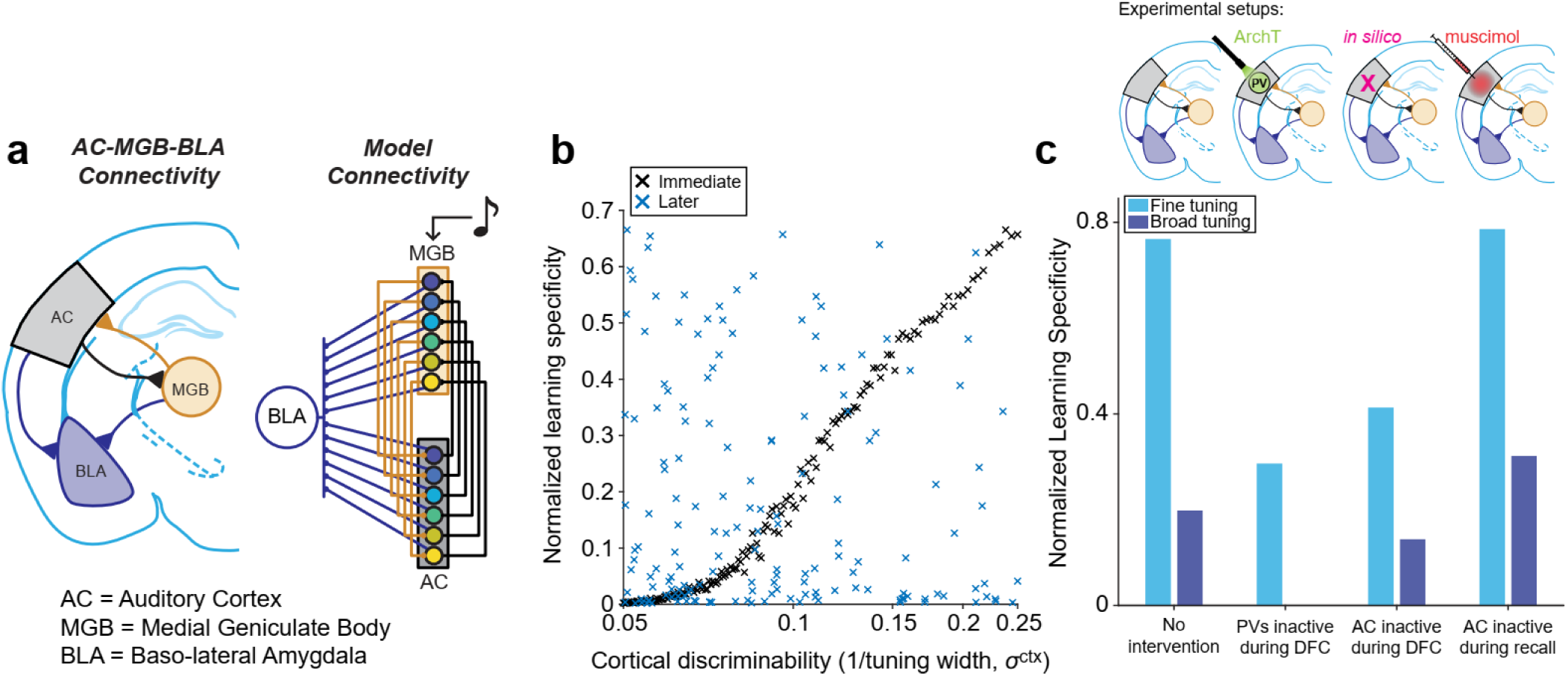
A learning model reconciles present and past findings. **(a)** (left) connectivity between auditory cortex (AC, gray), medial geniculate body (MGB, orange) and basolateral amygdala (BLA, blue). (Right) Model connectivity. MGB receives auditory input and provides input to AC (orange lines), and both MGB and AC provide inputs to BLA (blue lines). AC feeds back to MGB (black lines). Colored circles represent neurons tuned to different, overlapping frequency ranges. **(b)** Normalized learning specificity output from the model with varying levels of AC discriminability, achieved by changing the frequency tuning overlap between the neurons in the AC population, *σ^ctx^*. Learning specificity was measured at two time points, immediately after DFC (black) and 10^4^ time-steps later (blue). **(c)** Normalized learning specificity at two AC discriminability levels; fine (light blue) and broad (dark blue) tuning. Results are shown for learning specificity with no interventions, when inhibition is reduced in AC during DFC (analogue of when ArchT-transfected PV interneurons in AC are inactivated by optogenetics during DFC), when AC is inactivated during DFC (in the model), and when AC is inactivated during memory recall (analogue of an injection of muscimol during memory recall; PV = parvalbumin positive interneurons, ArchT = Archaerhodopsin-T).

First, we first tested whether broad tuning in AC (low neuronal discriminability between CS+ and CS-) during conditioning produced more generalized freezing than sharp tuning (high neuronal discriminability). We found that increased overlap in frequency tuning in AC neurons, without changing the tuning of MGB neurons, drove more generalized freezing responses (Fig 7b, S12). This is due to the fact that, when AC was broadly tuned, CS+ tone activated AC neurons not only responded to the CS+ frequency but also to other frequencies, such as the CS-, albeit to a lesser extent. After learning, this resulted in strong AC to BLA synaptic weights that are not specific to CS+. MGB is narrowly tuned in our model, but the weights from MGB to BLA were also strengthed in a non-specific fashion because AC projects back to MGB. Therefore, CS+ also activated non-specific neurons in MGB concurrently with the foot-shock. These results support the present findings (Fig 2, 3). Drift in the tuning properties of the neurons in the model led to the correlation between learning specificity and tuning width decreasing over time since conditioning, consistent with our finding that neural acticity post-DFC no longer predicts learning specificity (Fig 4). Second, we examined the effects of decreasing inhibition in the AC population during conditioning (Fig 7c, S13). Decreasing inhibition resulted in an increased overlap in frequency responses in the AC population, which in turn led to increased generalization, supporting previous findings and providing a mechanism^12,33^. Third, we tested the effects of inactivating AC during conditioning we found that learning specificity was reduced, consistent with the hypothesis that AC affects tone discrimination *during* DFC (Fig 7c, S14). Finally, we tested whether inactivating the auditory cortex *following* conditioning had an effect on freezing responses (Fig 7c, S15). Consistent with previous findings^3,11^, we did not observe a change in fear generalization following AC inactivation. The broad or narrow tuning of AC neurons allowed for the synapses from MGB to BLA to be strengthened either narrowly or broadly during conditioning. Therefore, with suppression of AC during memory recall, the specialized versus generalized learning was preserved.

Combined, the model demonstrates that a simple anatomically consistent circuit supports multiple aspects of cortical control of fear conditioning identified here and in previous studies.

## Discussion

Our results identify the role of the auditory cortex in differential fear learning: (1) Prior to fear learning, neuronal responses in AC shape fear learning specificity (Fig 2 & 3); (2) Following differential fear conditioning, neuronal response transformations are not correlated with fear learning specificity (Fig 5, Fig S8), and therefore the auditory cortex does not encode auditory differential fear memory; (3) Neuronal activity in AC post-DFC does not correlate with freezing behavior (Fig 4); (4) A simple model of the auditory nuclei and the basolateral amygdala could account for our results as well as a number of previous findings (Fig 7).

Our finding that the neuronal activity prior to fear conditioning predicted specialization of fear learning provides a mechanism for the role of AC in differential fear memory acquisition ^10–12,14,33^. Specifically, inactivation of inhibitory neurons in the AC during fear conditioning led to increased generalization of fear learning with pure tones ^12^. Suppressing inhibitory neurons in the AC led to a decrease in Fisher information, which reflects the certainty about a stimulus in neuronal representation^33^. This change would likely result in a decrease in neuronal discriminability between the dangerous and safe tones in the AC, and therefore drive a increase in fear generalization, as demonstrated by our model (Fig 7). Our results provide the link between optogenetic inactivation of interneurons in AC leading to increased fear generalization, and to increased frequency tuning width^12^, which decreases neuronal discriminability.

By using two-photon imaging to record from the same neurons over the course of differential fear conditioning, we were able to compute changes in both absolute and relative neuronal activity of a large number of identified neurons, a feat not normally achievable with electrophysiology^17,19^. Previous work found that changes in neuronal responses to the dangerous and safe stimuli after differential fear conditioning amplified the difference between the responses^17,19^. This change was proposed to represent fear memory^15,19,29^. We identified similar transformations in the *normalized* response functions of neurons that were tracked pre- to post-conditioning, we found an increased relative response to the CS+. However, these changes did not correlate with freezing behavior suggesting that the neuronal code in the AC after fear conditioning does not reflect differential fear memory. Indeed, a number of studies found that inactivating the auditory cortex *after* fear conditioning with pure tones does not affect fear memory retrieval^3,11^ (but see^14^). Combined, our results restrict the role of auditory cortex in fear conditioning to pure tone differential fear memory acquisition, but not retrieval.

If the increase in normalized response at CS+ is not related to fear memory, then why is there an increase in response? It could be reflective of increased attention caused by presentation of the CS+ and that the discrimination of the CS stimuli is unaffected by this effect^34^. Furthermore, changes in frequency map organization do not necessarily relate to changes in behavioral frequency discrimination of pure tones^35,36^, thus over-representation of the CS+ could be induced by learning but not necessary for discrimination learning.

To locate our findings with previous work, we implemented a simple, anatomically accurate^37,38^ model with connections from auditory nuclei to the basolateral amygdala (Fig 7). The model demonstrated that (1) neuronal activity in cortex can predict subsequent learning specificity; that (2) inactivation of PV interneurons in AC during DFC leads to increased generalization^12^, and that (3) the auditory cortex is not necessary for differential fear memory retrieval^3,11^ and (4) that discrimination is still possible with AC inactive during conditioning but learning specificity is reduced. The model proposes that either MGB or AC or a combination of both can induce auditory fear memory through the strengthening of connections in the amygdala. We propose that feedback from auditory cortex to the MGB contributes to discrimination of perceptually similar pure tone stimuli *during* DFC by controlling stimulus discrimination in the MGB, this may or may not be a direct projection neuroanatomically^37,39,40^. Random drift accounts for the lack of correlation between neuronal tuning and learning specificity after conditioning^27^. Future studies need to explore the role of the MGB and specific projections between AC, MGB and BLA in fear learning and memory. It is likely that such an important behavioral modification as fear has redundant pathways to obtain the same behavioral outcomes^11,41–43^.

Our results relied on tracking the neuronal responses in all transfected neurons in AC without distinguishing between different neuronal subtypes. Previous studies found that a specific class of inhibitory neuron increases activity with presentation of repeated tones^18,26^. It is therefore plausible that our results include a subset of neurons that function differently during fear conditioning but which we are unable to identify due to lack of selective labelling. Furthermore, we restricted our recordings to layers 2 and 3 of the auditory cortex, and it is possible our results overlook more specific changes in the thalamo-recipient layers of the cortex^44,45^. The complexity of transformations in the cortical microcircuit and between layers with learning can be explored further^46–49^.

The results of the study may be restricted to pure tone stimuli. We chose pure tone stimuli because these stimuli provide a well-defined axis (frequency) along which to vary stimulus discriminability and there is strong evidence to suggest auditory cortex modulates discrimination of pure tones^35,50,51^. Furthermore, in human subjects, AC encodes threat during DFC for pure tone stimuli ^21^. Our prior work has established that large frequency separation between CS+ and CS- results in uniform specificity of the fear response among subjects, whereas smaller frequency separation, such as the one used here, provides for a gradient of specificity across subjects^3,12^. Other studies have found that AC is not behaviorally relevant for discrimination between pure tones separated by large frequency distances^11,13^. However, when the frequencies were brought closer together, then manipulation of AC activity did affect behavior^13^. Therefore, it is unclear whether recent conclusions that AC is involved in processing of more complex stimuli and not pure tones are due to differences in complexity of the stimulus, or to the degree to which AC can discriminate these stimuli. Furthermore, the FM sweeps used in these studies are not necessarily more complex than pure-tones for AC processing. Indeed, neurons in the inferior colliculus, which is two synapses earlier than AC, differentiate between FM sweeps, e.g.^52^. Ultimately, the relevant aspect of the present study was the ability to measure how well neuronal ensembles differentiate between two stimuli. We achieved this by bringing CS+ and CS- close together in frequency, and we found that neuronal discriminability of the stimuli differs across mice and correlates with behavioral discriminability prior to DFC. We would not expect this result were the stimuli not relevant for AC. Furthermore, inactivation of AC during conditioning in the model led to a decrease in learning specificity (Fig 7). Future studies will dissect to what extent the differences in neuronal codes in AC shape differential fear learning of more complex and natural sounds and its role in other forms of learning^13,36,53,54^.

Our results may be applicable to understanding anxiety disorders. An extreme example of fear generalization is realized in PTSD^55^. Here we find that the present state of each individual brain, in terms of neuronal discrimination of stimuli, is predictive of the future generalization of fear in the subject. This suggests that a way to prevent generalization of dangerous and safe sounds is to improve neuronal discrimination of potentially threatening stimuli^56–59^. Further work in this area can lead to a deeper understanding how genetic and social factors, as well early life experiences, shape the role of sensory cortex in this common and devastating disorder^7,58^.

We identified a neuronal correlate for inter-individual differences in learning specificity. We found that the mammalian sensory cortex plays key role in stimulus discrimination during, but not following, differential fear conditioning. These results reconcile several previous findings and suggest that the role of sensory cortex is more complex than previously thought. Investigating the changes in the cortico-amygdala circuit during fear learning will pave way for new findings on the mechanisms of learning and memory.

## Acknowledgements

The authors thank Dr. Yale Cohen, Dr. Steve Eliades and Dr. Jay Gottfried on helpful discussions and comments on an earlier version of the manuscript. The authors also thank other members of the Geffen laboratory for their helpful advice. This work was supported by funding from the National Institute of Health grants R01DC015527, R01DC014479 and R01NS113241 to MNG.

## Author Contributions

Conceptualization, M.N.G. and K.C.W.; Methodology, K.C.W, M.N.G., C.F.A. and C.C.; Software, K.C.W.; Investigation, K.C.W. and K.O..; Formal Analysis, K.C.W. and M.N.G; Writing – Original Draft, K.C.W., C.C, M.N.G.; Writing – Review and Editing, K.C.W., M.N.G. and C.C.; Funding Acquisition, M.N.G.; Resources, M.N.G., K.O. and K.C.W.; Supervision, M.N.G.

## Online methods

### Mice

All experimental procedures were in accordance with NIH guidelines and approved by the IACUC at the University of Pennsylvania. Mice were acquired from Jackson Laboratories (19 male, 9 female; age 12 weeks, PV-Cre (4) [Stock No: 017320], CamKII-Cre mice (1) [Stock No: 005359] or Cdh-23 mice (23) [Stock No: 018399]) and were housed in a room with a reversed light cycle. Experiments were carried out during the dark period. Mice were housed individually after the cranial window implant. 19 mice (13 males, 6 females) were in the conditioning group and 9 mice (6 males, 3 females) were in the *pseudo*-conditioned control group.

The Auditory Brainstem Response (ABR) to tone pips (4 – 32 kHz, 10 – 80 dB SPL) was acquired before or at the end of the experiment, when possible, in order to confirm that mice had thresholds for the stimuli at or below the presentation level (Fig S16). Mice with ABRs >70 dB were excluded from the study (*N* = 2).

Euthanasia procedures were consistent with the recommendations of the American Veterinary Medical Association (AVMA) Guidelines on Euthanasia.

### Surgical procedures

Mice were implanted with cranial windows over auditory cortex. Mean age of cranial window implant: 9.6 weeks [6.3 – 13.0 weeks]. Briefly, mice were anaesthetized with 1.5 – 3% isoflurane and a 3-mm circular craniotomy was performed over the left auditory cortex (stereotaxic coordinates) using a 3-mm biopsy punch centered over the stereotaxic coordinates of A1 (70% of the distance between bregma and lambda, 4.3 mm lateral to the midline). An adeno-associated virus (AAV) vector encoding the calcium indicator GCaMP6s or GCaMP6m (AAV1.Syn.GCaMP6s.WPRE.SV40 or AAV1.Syn.GCaMP6m.WPRE.SV40, UPENN vector core) was injected (750 nl, ∼1.89 x 10^-12^ genome copies·ml^-1^) at a 750µm depth from the surface of the brain at 60 nl min^-1^ for expression in layer 2/3 neurons in A1. 3 injections were made at the same lateral distance but separated by 0.5 mm in the anterior-posterior direction or 5 injections were made spread across the window (0.3 – 0.5 mm apart). The injection needle was left in place for 10 mins after the injection was complete before retraction. Injections were made using a pump (Pump 11 Elite, Harvard Apparatus, USA) and needles were pulled (P-97 Puller, Sutter Instruments, USA) glass pipettes (Harvard Apparatus, USA) with tip openings of 30 – 50 µm. After injection, a circular 3-mm diameter glass coverslip (size 0 or 1, Warner Instruments) was placed in the craniotomy and fixed in place using a mix of cyanoacrylate glue and dental cement. A custom-made stainless-steel head-plate (eMachine Shop) was fixed to the skull using C&B Metabond dental cement (Parkell). The implant was further secured using black dental cement. Mice were allowed to recover for 3 days post-surgery.

### Behavioral training and testing

Mice underwent a minimum of 4 imaging sessions (range: 4 – 11) prior to differential auditory fear conditioning (DFC). DFC and subsequent fear retrieval testing took place in two different contexts (A and B, discussed below). Before and after each conditioning or retrieval, we cleaned the conditioning and testing chambers with either detergent (retrieval chamber) or 70% ethanol (conditioning chamber). We recorded a video of the mouse in the testing chamber using FreezeFrame 3 software (Coulbourn) at 3.75 Hz; the subsequent movement index (mean grayscale values of frame (*n*+1) minus the preceding frame (*n*)) was exported and analyzed offline using MATLAB. The threshold of movement was defined as the 12.5^th^ percentile of the values from each session. The mouse was considered to be freezing if the movement index was below the threshold; the measure of freezing was expressed as a percentage of time spent freezing during stimulus presentation and for baseline during the 30s prior to stimulus onset.

Stimuli were generated using FreezeFrame 3 and presented at 70 dB SPL from an electrostatic speaker (ES-1, TDT) mounted above the animal. DFC took place in context A (Fig 1). Stimuli were 30 s in duration and were either a continuous pure tone (4 mice) or pulsed pure tones (500 ms duration at 1 Hz). The CS+ (15 kHz) was paired with a foot-shock (1 s, direct current, 0.7 mA, 10 pairings, inter-trial interval: 50 – 200 s) delivered through the floor of context A (by precision animal shocker, Coulbourn). The foot-shock either co-terminated with the continuous tone or the onset coincided with the final tone pulse of the CS+ stimuli. The CS- (11.4 kHz) was presented after each CS+-foot-shock pairing but was not reinforced (10 presentations, inter-trial interval: 20 – 180 s). Fear memory retrieval sessions in context B followed each two-photon imaging session after conditioning. The CS+ and CS- were presented 4 times (30 s duration, interleaved, inter-trial interval: 30 – 180 s). For 4 mice, longer continuous presentations of the CS+ and CS- were presented (either 120 s, 1 mouse, or 60 s, 3 mice), for these mice, trials were divided into 4 equal durations and treated as above. In *pseudo*-conditioning, the foot-shocks were presented interleaved between the stimuli in periods of silence. Baseline freezing consisted of an equal time of silence prior to tone onset.

Conditioned mice that did not freeze either to CS+ or CS- were removed from subsequent analysis (two-way ANOVA for each mouse on freezing scores to CS+, CS- and baseline from all retrieval sessions (16 trials for each CS and 32 trials for baseline). Stimulus (CS+/CS-) and baseline (stimulus/no stimulus) were the independent variables. Learners were defined as those with significant effect of baseline or baseline*stimulus, *p* < .05). 6 mice (5 males, 1 female) were excluded from the study, leaving 15 conditioned mice (10 males, 5 females).

For each mouse the learning specificity (*LS*, Equation 1 ^3^) was calculated as:

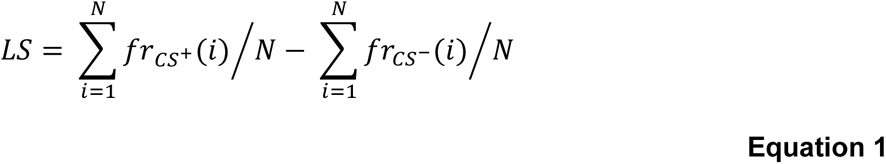

Where *i* is the trial index, 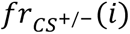 is the fraction of time spent freezing during trial *i* in the CS+/- condition, respectively, and *N* is the number of trials per condition.

### Calcium imaging procedure and acoustic stimuli

All imaging sessions were carried out inside a single-walled acoustic isolation booth (Industrial Acoustics). Mice were placed in the imaging setup, and the head plate was secured to a custom base (eMachine Shop) serving to immobilize the head. Mice were gradually habituated to head-fixing over 3 – 5 days, 3 – 4 weeks after surgery and before imaging commenced. Imaging took place in mice aged at the end of experiments 19.6 ± 2.5 weeks ± sem.

We recorded changes in fluorescence of GCaMP6s/m caused by fluctuations in calcium concentration in transfected neurons of awake, head-fixed mice, using two-photon microscopy (Ultima *in vivo* multiphoton microscope, Bruker). We used a 16X Nikon objective with 0.8 numerical aperture (Thorlabs, N16XLWD-PF). The laser (940 nm, Chameleon Ti-Sapphire) power at the brain surface was kept below 30 mW. Recordings were made at 512 x 512 pixels and 13-bit resolution at ∼30 frames per second.

Stimuli were generated at a sampling rate of 400 kHz using MATLAB (MathWorks, USA) and consisted of 100-ms long tone pips in the 5−32-kHz frequency range presented at 60 – 80 dB SPL. In a single recording session, each frequency was repeated 15 – 30 times in a *pseudo*-random order with a 4-s inter-stimulus interval.

### Cell tracking across imaging sessions

We imaged the activity from the same cells over 15 days in layers 2/3 of auditory cortex, using blood vessel architecture, depth from the surface, and the shape of cells to return to the same imaging site. To identify ROIs across imaging sessions that corresponded to the same cell, the maximum-projection fluorescence images from each day were registered by transforming the coordinates of landmarks present in both images in MATLAB (2017a) using the *fitgeotrans* function. The transformation was applied to ROIs from the second imaging session to match the first – all subsequent sessions were aligned to the first imaging session. We next calculated the distance between all the pairs of centroids (mean x-y position of each ROI) across the two sessions; ROIs from the two sessions were then automatically registered as the same cell based on the nearest centroid. We then manually checked the shape and position of the ROIs for any pairs that had duplicate matches, <80% ROI overlap, or a larger than average distance between the centroid locations (>2 standard deviations). ROIs which were not matched to any earlier ROIs were counted as new cells. This process was repeated for subsequent sessions, registering the imaging field to the first session, and comparing the ROIs to the cumulative ROIs from previous sessions. A final manual inspection of all the unique ROIs was performed after all the imaging sessions were registered. ROIs that overlapped with each other extensively were excluded from the dataset since it was unclear whether they were the same or different cells. Examples of tracked cells and aligned ROIs are shown in Fig S1.

### Data analysis and statistical procedures

Publicly available toolboxes^22^ running on MATLAB were used to register the two-photon images, select regions of interest (ROI), and estimate neuropil contamination, resulting in a neuropil-corrected fluorescence trace (*F*) for each neuron (*F* = trace - (neuropil*0.7)). This trace was low pass filtered (filter cut off at 7.5 Hz) to remove high frequency noise. From this filtered trace, we calculated the mean baseline fluorescence (*F_baseline_*) and standard deviation of the baseline (*F_std_*) over the one second prior to tone onset, and then determined the change in fluorescence over time relative to the mean baseline fluorescence (*ΔF = F - F_baseline_*) for each sound presentation. We then divided *ΔF* by *F_std_*, effectively calculating the z-score of the fluorescence response relative to the baseline (*ΔF/F_std_*) for each sound presentation.

The response to each tone was defined as the mean *ΔF/F_std_* over 2 seconds following tone onset. Neurons were deemed sound responsive if at least one of the frequency responses was different from zero (*t*-test, *p* < 0.05, corrected for multiple comparisons using false discovery rate ^60,61^). The frequency response function was defined as the mean response to each tone frequency across repeats. Best frequency was defined as the frequency with the highest mean response. Sparseness (*S*, Equation 2^30,31^) was used to estimate the sharpness of response functions, with 1 being very sharply tuned and 0 being an equal response to each tone frequency:

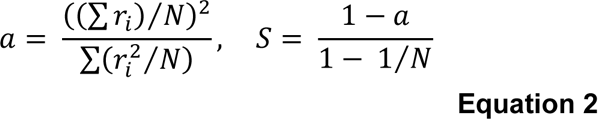

where *r_i_* is the mean response to the frequency *i* and *N* is the total number of frequencies tested.

The Z-scored difference between responses to CS+ and CS- (Z_diff_, Equation 3) was calculated for each neuron using the following equation:

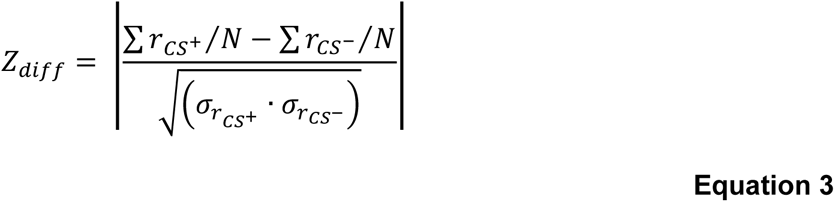

where *r_CS+/CS-_* is the single trial mean responses (mean *ΔF/F_std_* over 2 s post-stimulus onset) to CS+ and CS- respectively, *N* is the number of repeats of each stimulus and *σ* is the standard deviation of mean responses. The Z_diff_ score was considered significant if the actual Z_diff_ was larger than the 95^th^ percentile of the distribution of Z_diff_ scores calculated with shuffled the CS+/CS- response labels 250 times. For mice not tested under the two-photon directly with the CS+ or CS-, the data were linearly interpolated to estimate responses at CS- and CS+. We used average Z_diff_ across pre-DFC sessions of learner mice to test whether there was a difference between using GCaMP6s (6/23) and GCaMP6m (17/23). We found no difference (unpaired t-test, *t*(21) = 1.04, *p* = .309) between the mean Z_diff_ scores of the two groups of mice and thus we have analyzed them together.

For fitting the Support Vector Machine, we used MATLAB’s *fitcsvm* function with a Linear kernel and 10-fold cross-validation to predict the learning specificity based on the standardized single-trial population responses (mean *ΔF/F_std_* over 2 s post-stimulus onset for each neuron).

We calculated the confidence intervals of correlations using a bootstrap procedure, resampling, with replacement, the data 1000 times, and computing the Pearson’s correlation between the resampled data. We defined the 95% confidence limits of the correlation coefficient (*ρ*) as the 2.5^th^ and 97.5^th^ percentiles of the resulting distribution of correlation coefficients. In order to assess whether two correlations were significantly different from one another we subtracted the bootstrapped *ρ* distributions of each dataset from one another, the change in *ρ* was considered significant if 95% CI of the difference-distribution did not overlap with zero.

To compare results between testing groups (conditioned/*pseudo*-conditioned) we used two-way repeated measures ANOVAs, linear regressions, and linear mixed-effects models with the relevant variables (see Table S1-3).

For mice that were not tested at 11.4 and 15 kHz under the two-photon microscope (4 conditioned mice) responses were linearly interpolated from the frequency response functions pre- and post-DFC. For cells present in more than one session either pre- or post-DFC, the frequency response curves from each session were averaged and the changes in response were assessed from the mean across pre- and post-DFC sessions. Sparseness and best frequency were calculated from the mean responses across sessions. Z_diff_ scores were also averaged across neurons responsive in multiple sessions pre- and post-DFC. For comparing the fluorescence traces of responses (Fig S7d-g), for the 4 mice not tested directly at CS+ and CS-, the nearest frequencies were used.

### Confirming anatomical location of recording

Upon conclusion of the imaging sessions, we removed the windows of the mice and injected a red fluorescent marker (Red Retrobeads, CTB or AAV5.CAG.hChR2(H134R)-mCherry.WPRE.SV40 (mCherry)) into the site of imaging as identified by blood vessel patterns. Briefly, we anaesthetized mice with 1.5 – 3% isoflurane and used a drill (Dremel) to remove the dental cement holding the window in place. We removed the glass window and injected the red marker into the imaging site (Red Retrobeads: 250 nl, CTB: 500 nl (0.5%), mCherry: 500 nl) using a glass pipette (tip diameter: 40 – 50 µm) at 60 nl min^-1^. Following the injection, we covered the exposed brain with silicon (Kwik-Sil, World Precision Instruments) and then coated it with dental cement. After allowing time for retrograde transport (retrobeads and CTB: 1 week) or viral transfection and expression (mCherry: 3 weeks) mice were deeply anesthetized with a mixture of Dexmedatomidine (3 mg/kg) and Ketamine (300 mg/kg) and brains were extracted following perfusion in 0.01 M phosphate buffer pH 7.4 (PBS) and 4% paraformaldehyde (PFA). They were further fixed in PFA overnight and cryopreserved in 30% sucrose solution for 2 days before slicing. The location of imaging was confirmed through fluorescent imaging (Fig S2). For Retrobeads and CTB, the injection site was clear as a very bright injection site, for mCherry, expression levels were measured across the AC and the site of imaging was assumed to be the section with the strongest expression/brightest red. The identified sections were cross-referenced with the Allen Institute Mouse Brain Atlas using freely available software ^62^.

### Model

#### Neuron model and network

We simulated cortical neuronal populations, MGB populations and a BLA neuronal population in a rate-based description of neuronal activity. We simulated *N* = 10 MGB populations. Each MGB population receives *N* = 10 inputs *x_i_^MGB^*, *i* = 1..N. To model the fact that neighboring inputs are correlated, we generated the inputs *x_i_* assuming that they each have a similar tuning to stimuli. These stimuli were modeled as 10 time-dependent activities *s_j_*(*t*) (which corresponded to a sound amplitude at a given frequency, *j*). The activity of input *i* was calculated by a sum of the stimulus channels, weighted with tuning strengths 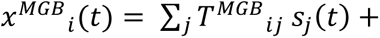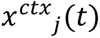. The input tuning was Gaussian: 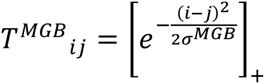 for *i* and *j* going from 1 to 10. [. ]_+_ means that negative values are set to zeros. The term *x^cxt^* corresponds to the direct cortical feedback. The parameter *σ^MGB^* regulated how broad the population response is to the sound. In the model, we assumed that MGB neuronal populations always have a small overlap in neuronal responses (*σ^MGB^* = 0.8).

Similarly, we simulated *N* = 10 cortical populations as 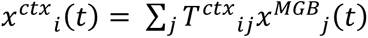. The input tuning was also Gaussian: 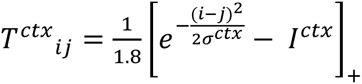 for *i* and *j* from 1 to 10. *I^ctx^* = 0.9 was a broad inhibitory term.

In the simulations, we tested for two different values of initial *σ^ctx^*; one corresponding to narrow tuning with a small overlap (*σ^ctx^* = 3), and one corresponding to broad tuning with a large overlap (*σ^ctx^* = 10). (Note that *σ^MGB^* = 0.8 was equivalent to *σ^ctx^* = 3 since we did not model MGB inhibition here, *I^MGB^* = 0). To avoid boundary effects, we had a circular boundary condition of the 10 inputs, meaning that input 1 and input 10 are neighbors. We also assumed that the tuning *σ^ctx^* would drift over time. Specifically, at every time step, we added a uniform random noise between -0.25 and 0.25 to *σ^ctx^*. *σ^ctx^* was bounded between 4 and 20.

Finally, we simulated one population in the BLA. It received inputs from both cortical and MGB populations, i.e., *y* = *w^MGB^ x^MGB^* + *w^ctx^ c^ctx^*, where *w^MGB^* are the weights from MGB neurons to the BLA neurons, and *w^ctx^* are the weights from cortical neurons to the BLA. Normalized freezing response was computed as the activity after the fear conditioning paradigm (see below) normalized by the maximal activity (i.e., when the weights are all 1).

#### Modelling fear conditioning paradigm and interventions

During the fear conditioning training to simulate a CS- tone, we set (channel number 6) *s*_6_ = 1, all the other inputs to zero, and a CS+ we set (channel number 3) *s*_3_ = 1, all the other inputs to zero. In addition, we paired it with a shock (*e* = 1 if there is a shock, *e* = 0 otherwise). The synaptic weights were plastic under the following rules: 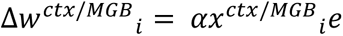, where α = 0.1 is the learning rate. This is analogous to the standard Delta rule. The weights were bound between 0 and 1 and are initialized at 0.1. We simulated the fear conditioning for 10 time-steps [arbitrary time] and spontaneous dynamics with tuning *σ^ctx^* drift for another 10000 time steps. To simulate optogenetic inactivation of PV neurons in AC ^12^, which decreases inhibition in AC, we lowered inhibition in AC by setting *I^ctx^* = 0.45 (half the ‘normal’ level), the maximum freezing was computed with the original inhibitory term intact (*I^ctx^* = 0.9). To simulate pharmacological inactivation of AC during memory recall (after learning), we tested the behavior of the model with AC inactivation by setting 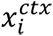 = 0 during the BLA simulation protocol.

### References

Recent work in several fields of science has identified a bias in citation practices such that papers from women and other minority scholars are under-cited relative to the number of such papers in the field ^63–67^. Here we sought to proactively consider choosing references that reflect the diversity of the field in thought, form of contribution, gender, race, ethnicity, and other factors. First, we obtained the predicted gender of the first and last author of each reference by using databases that store the probability of a first name being carried by a woman ^67,68^. By this measure (and excluding self-citations to the first and last authors of our current paper), our references contain 12.21% woman(first)/woman(last), 3.15% man/woman, 27.0% woman/man, and 57.64% man/man. This method is limited in that a) names, pronouns, and social media profiles used to construct the databases may not, in every case, be indicative of gender identity and b) it cannot account for intersex, non-binary, or transgender people. Second, we obtained predicted racial/ethnic category of the first and last author of each reference by databases that store the probability of a first and last name being carried by an author of color ^69,70^. By this measure (and excluding self-citations), our references contain 10.02% author of color (first)/author of color(last), 14.31% white author/author of color, 14.22% author of color/white author, and 61.45% white author/white author. This method is limited in that a) names and Florida Voter Data to make the predictions may not be indicative of racial/ethnic identity, and b) it cannot account for Indigenous and mixed-race authors, or those who may face differential biases due to the ambiguous racialization or ethnicization of their names. We look forward to future work that could help us to better understand how to support equitable practices in science.

## Supplemental Information

**Figure S1:**
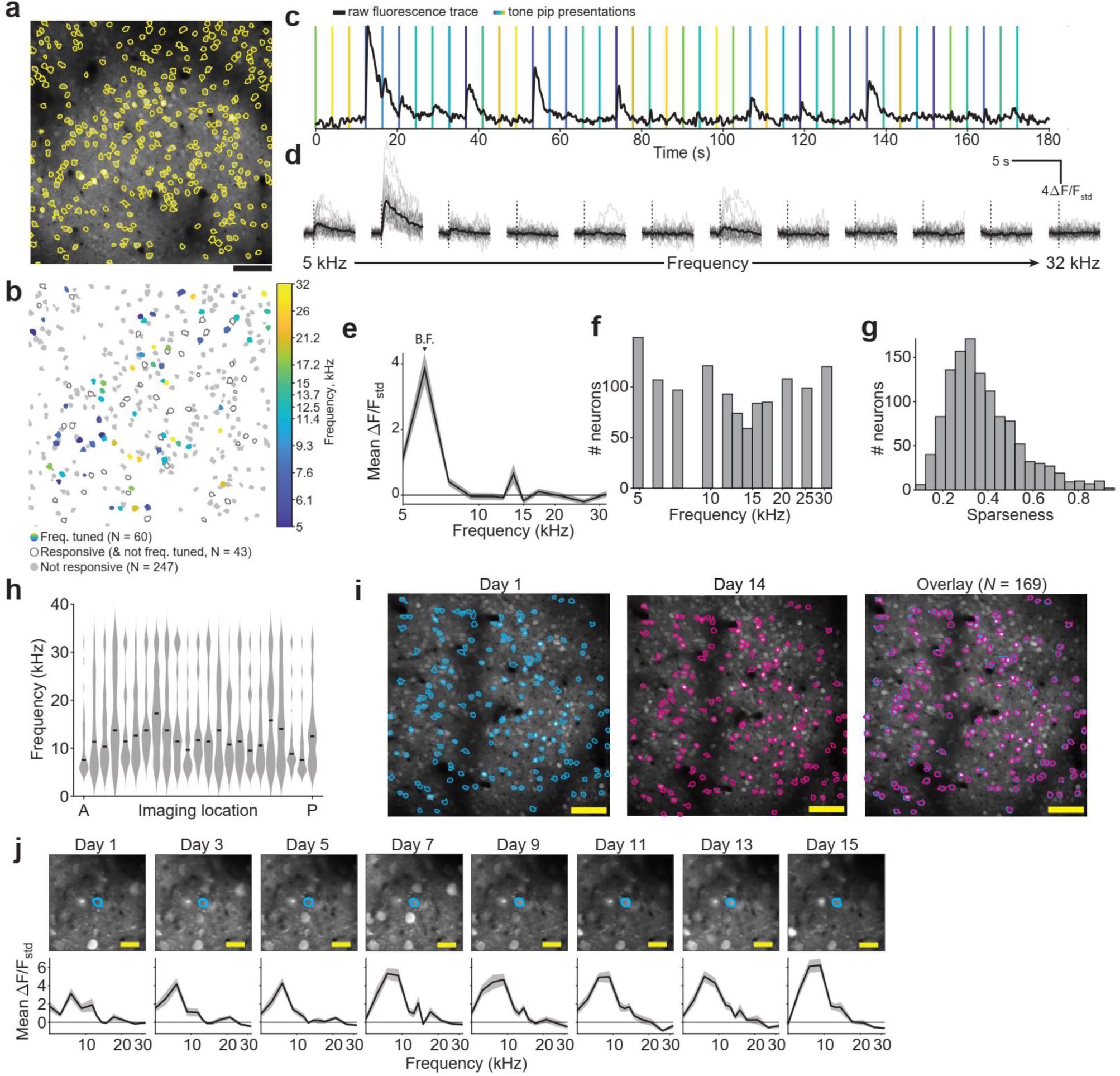
Longitudinal two-photon imaging tracks activity of neurons over weeks. (a) Two-photon imaging field of view with regions of interest corresponding to individual neurons (yellow outline, *N* = 350). Scale bar = 100 µm. (b) Cell outlines from a indicating cells not responsive to the stimuli (light gray), cells responsive to tones (dark gray, *t*-test against zero, *p* < .05, corrected for multiple comparisons using Holm–Bonferroni method) but not frequency tuned and frequency-tuned cells (colored according to frequency tuning, significantly responsive and one-way ANOVA, *p* < .05). Color bar indicates best frequency of each tuned neuron. (c) Part of a raw fluorescence trace (black) for an example frequency-tuned neuron with tone pip presentation times overlaid in color (vertical lines). The color of the vertical lines corresponds to the frequency of the tone pip presented – colors as in b. (d) Responses of neuron in c with single-trial responses (gray, *n* = 25 for each frequency) and the mean response (black). Dashed lines = tone pip onsets. (e) Mean response (from tone onset to 2 s after tone onset) across trials at each frequency of neuron in c-d. This neuron has a best frequency (B.F.) of 6.1 kHz. (f) Distribution of best frequencies of responsive cells recorded 24 hours pre-DFC (*N* = 1203, mice = 26). (g) Distribution of sparseness of responsive cells recorded in f. (h) Smoothed best frequency distributions ^71^ for each mouse pre-DFC ordered by anatomical location of the imaging window from anterior (A) to posterior (P). Black dashes indicate the median best frequency. (i) Field of view from two imaging sessions from the same mouse (maximum intensity projections), 15 days apart (left and middle) with ROIs tracked between the two sessions outlined in cyan and magenta. The right panel shows the ROIs from the two sessions overlaid. Scale bar = 100 µm. (j) Frequency responses of a representative cell over the 8 sessions of the experiment. Cell is shown outlined (cyan line). Scale bar = 25 µm. Error bars = standard error of the mean (sem)

**Figure S2:**
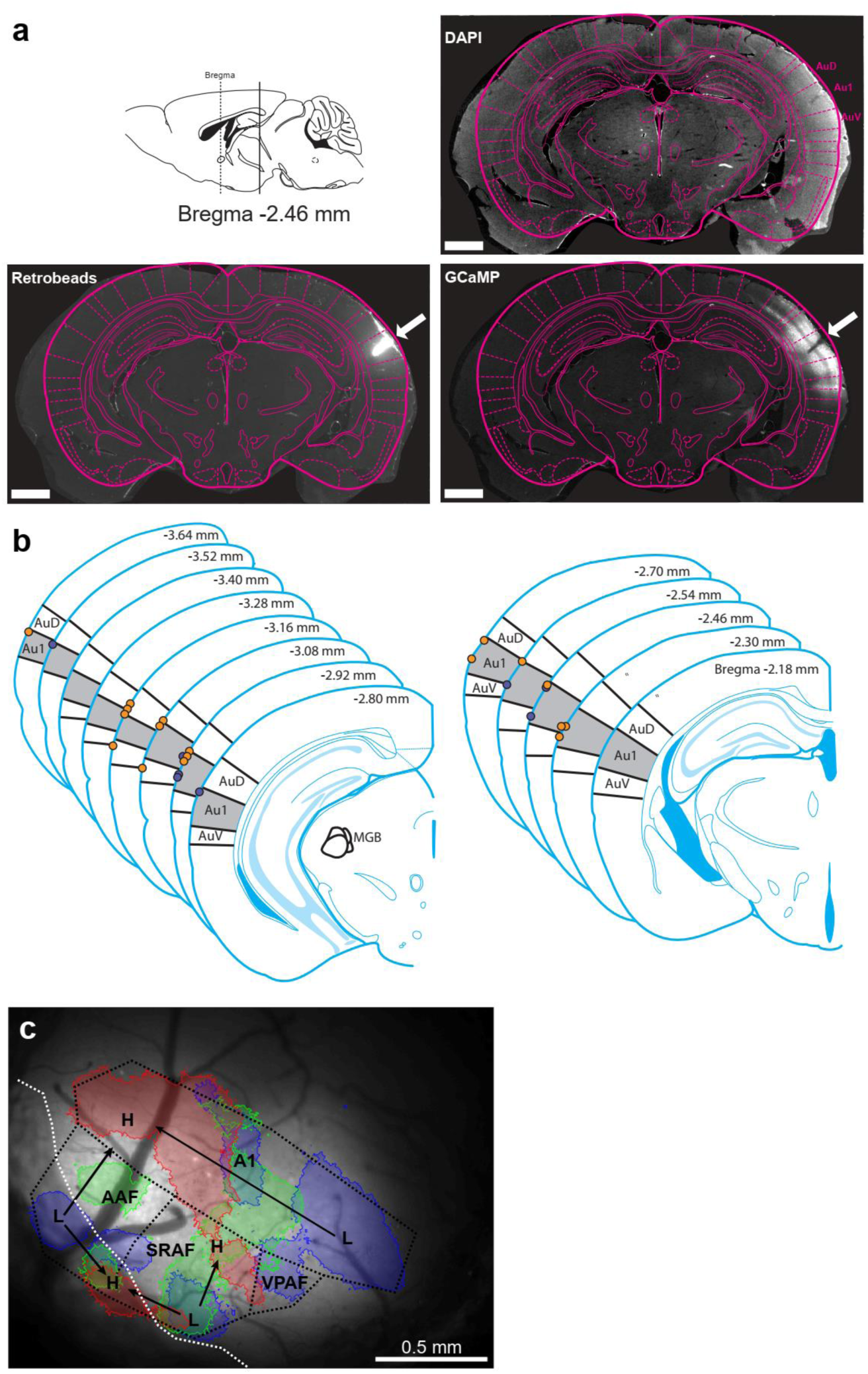
Location of imaging site example. **(a)** Upon completion of the experiments, the cranial window was removed, and a red indicator injected into the imaging site. The brains were fixed and sectioned at 40µm. We applied DAPI to reveal cell nuclei. This stain was used to align the section with the red indicator injection site (white arrows) with the mouse brain atlas (magenta outline). The red channel reveals the site of injection (Retrobeads) and is aligned with strong GCaMP6m/s expression (GCaMP). **(b)** Using the red injection sites as markers for the location of the imaging windows, the brain sections were aligned with the mouse brain atlas ^72^ to obtain the anterior-posterior location of the imaging windows relative to bregma and the field of auditory cortex imaged from. The center of the imaging field of view on the surface of the brain of conditioned (yellow) and pseudo-conditioned (blue) mice (N = 26/28) is indicated on the mouse brain atlas adapted from. The left stack of sections contains the MGB and the more anterior right-hand stack of sections does not contain the MGB Franklin & Paxinos, 3rd edition Mouse brain atlas. **(c)** For two mice, the red injection failed, and we used the widefield imaging to locate the imaging field of view. The figure shows an example widefield result. Thresholded responses to low (5 kHz, blue), medium (15 kHz, green) and high (30 kHz, red) tones of each pixel are indicated by the shaded regions. The pattern of responses allowed us to estimate the locations of the auditory fields using ^73^ as a guide. A1 – primary auditory cortex, VPAF – ventral posterior auditory field, SRAF – suprarhinal auditory field, AAF – anterior auditory field. Arrows indicate tonotopic gradients from low (L) to high (H) frequency.

**Figure S3:**
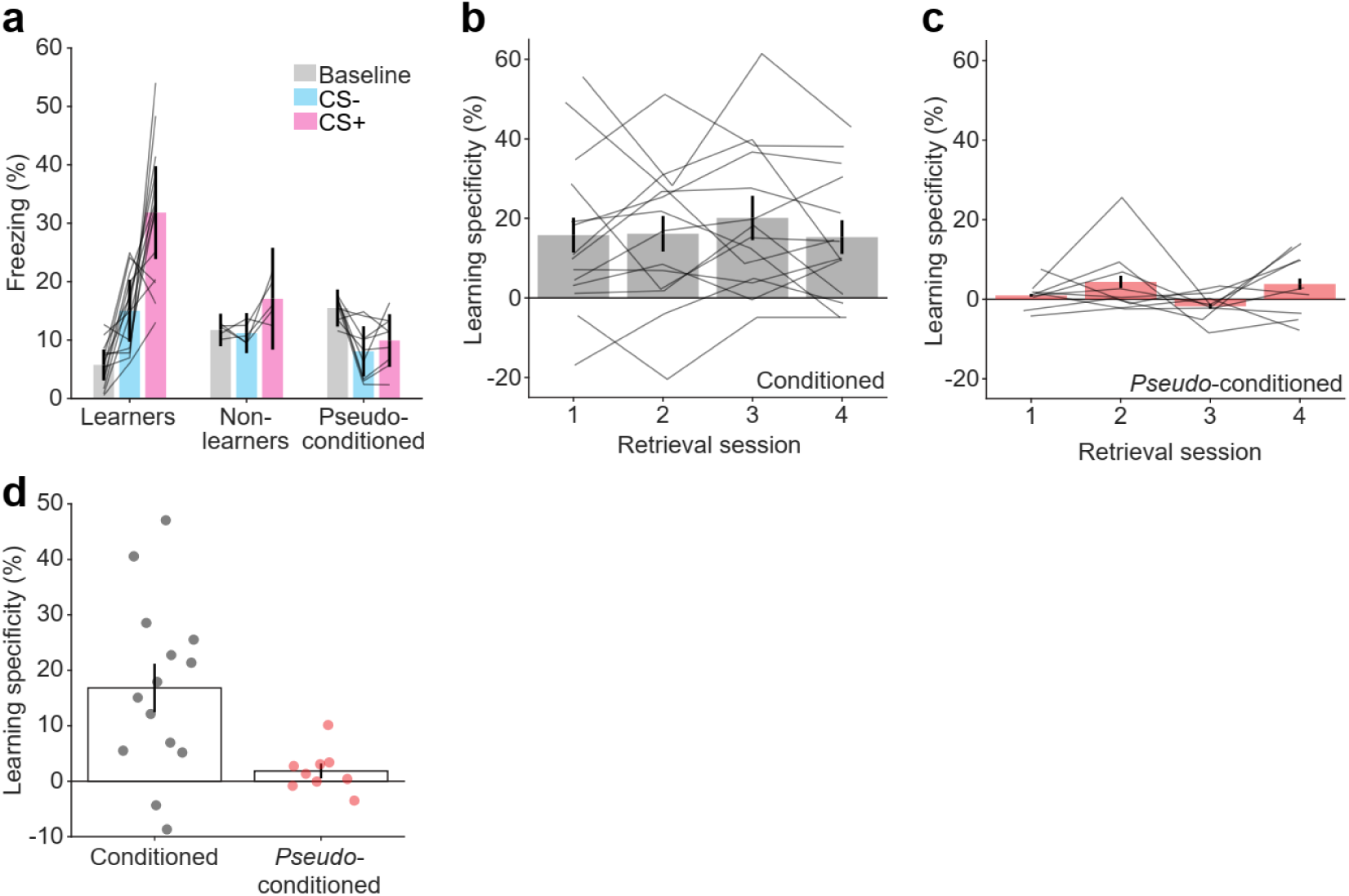
Mean freezing to CS+ and CS- and learning specificity across sessions. **(a)** Mean freezing across all 4 retrieval sessions at baseline (gray) and in response to CS+ (pink) and CS- (blue) for all mice. Learners (*N* = 14) showed a significant difference from baseline freezing in (baseline vs non-baseline) while non-learners (*N* = 5) did not (see Methods). Non-learners were subsequently excluded from further analysis. *Pseudo*-conditioned mice *N* = 9. Gray lines indicate individual mice. **(b)** Mean learning specificity in each retrieval session for conditioned mice. Gray lines show individual mice. **(c)** Same as **b** for *pseudo*-conditioned mice. **(d)** Mean learning specificity across all 4 retrieval sessions for conditioned (gray) and *pseudo*-conditioned mice (red). Circles show individual mice. Error bars in **a-d** show standard error of the mean.

**Figure S4:**
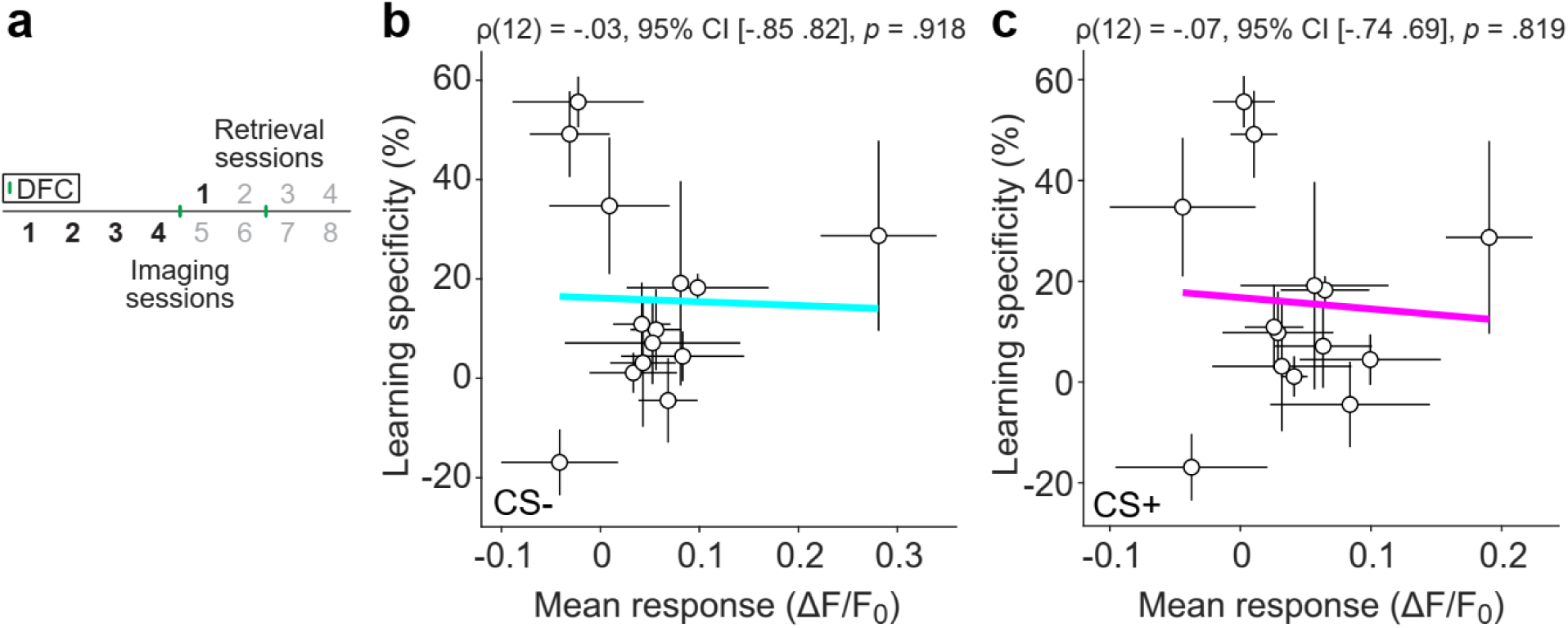
Mean response to CS+ or CS- does not predict learning specificity. **(a)** Responses to CS+ and CS- of responsive cells were averaged in each session and across the 4 pre-DFC imaging sessions (1-4). These mean responses were compared with the learning specificity from the retrieval session (1) 24 hours post-DFC. **(b)** Learning specificity of conditioned mice (*N* = 14) plotted against the mean response to CS- across the 4 pre-DFC imaging sessions. The line shows the linear best fit. Error bars represent standard error of the mean. Statistics: Spearman’s correlation. **(c)** Same as **b** but for responses to CS+.

**Figure S5:**
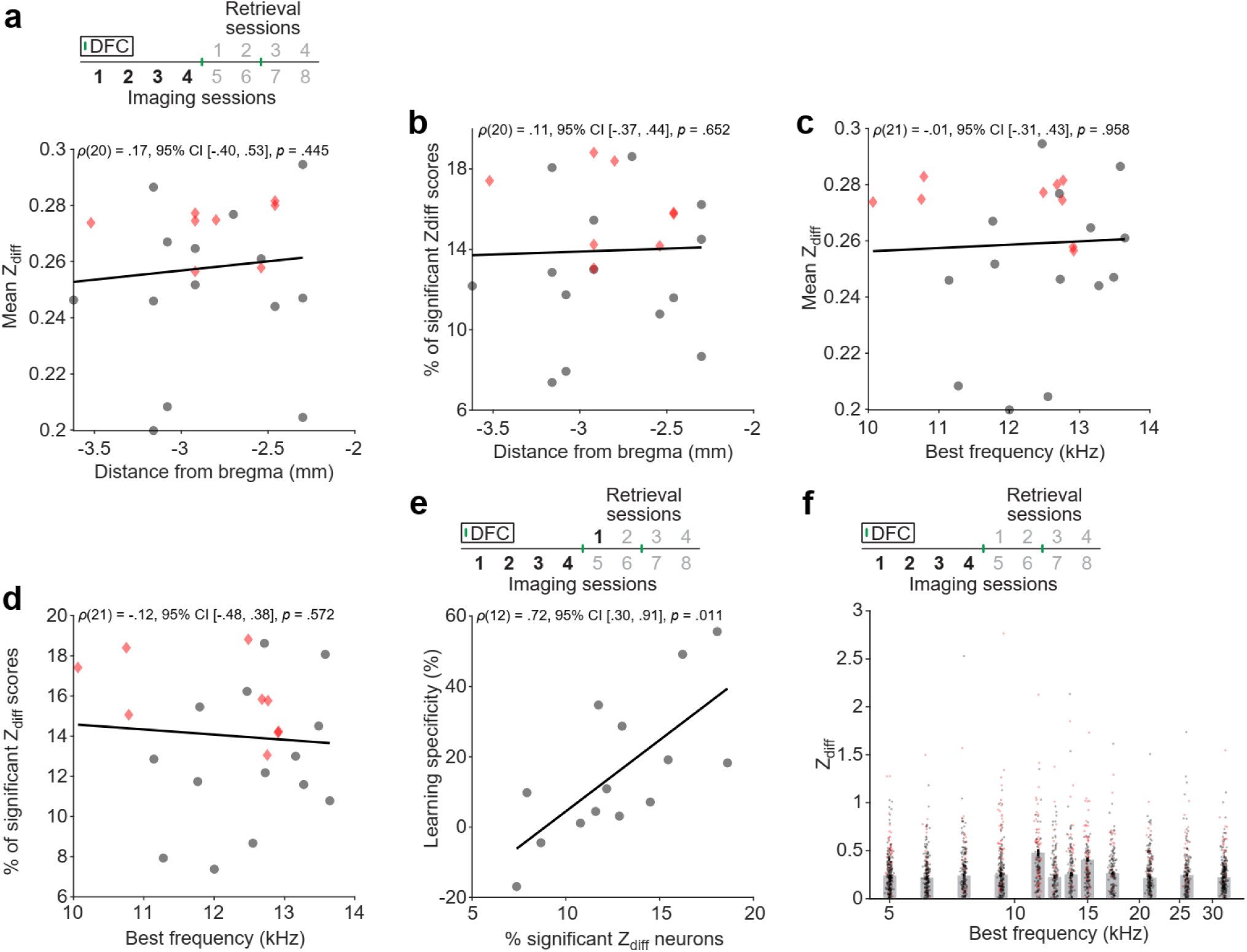
Relationship between Z_diff_ and best frequency distributions and anatomical location. **(a)** Number of responsive neurons with significant Z_diff_ and Z_diff_ of responsive neurons was averaged in each session and across sessions for each mouse. Relationship between the distance from bregma of the imaging field of view and the mean Z_diff_ scores of conditioned (black circles) and pseudo-conditioned mice (red diamonds) across pre-DFC sessions. **(c)** Relationship between the distance from bregma of the imaging field of view and the % of significant Z_diff_ scores across pre-DFC sessions. **(d)** Relationship between mean best frequency of responsive units in the imaging field of view and the mean Z_diff_ scores across pre-DFC sessions. **(e)** Relationship between mean best frequency of responsive units in the imaging field of view and the % of significant Z_diff_ scores across pre-DFC sessions**. (f)** Relationship between the mean % of significant Z_diff_ scores in the imaging field of view across pre-DFC sessions and the learning specificity from retrieval session 1 for conditioned mice. **(g)** Relationship between mean Z_diff_ across tracked pre-DFC sessions for each neuron and best frequency. Grey bars show median Z_diff_ ± sem. Statistics: Spearman’s rank correlation.

**Figure S6:**
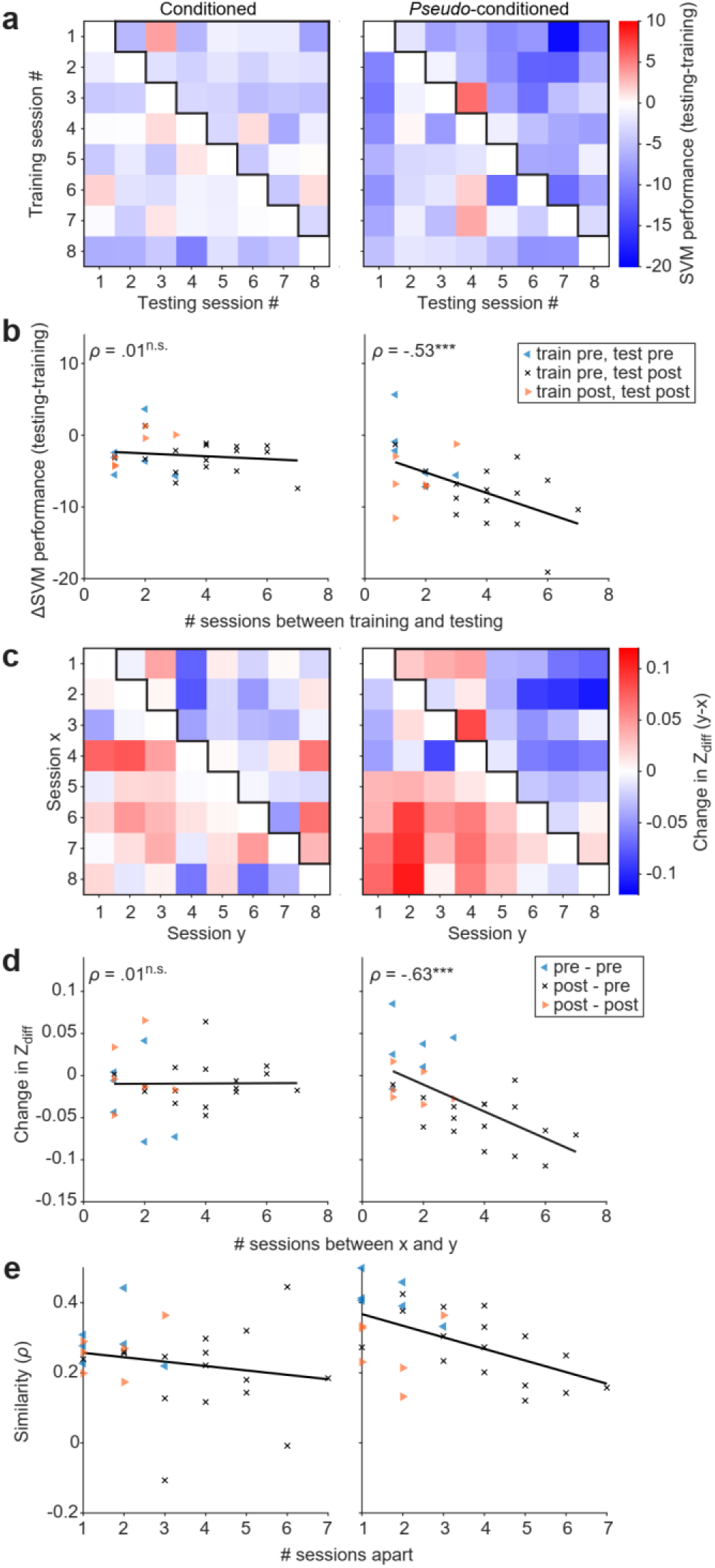
Changes in neuronal discrimination between imaging sessions. **(a)** The SVM was trained with data of neurons tracked between a pair of sessions from one of the sessions. The SVM was subsequently with data left out from that training set (10-fold cross validation) and with the same number of trials from the left-out neurons using the testing set. This was repeated across all pairs of imaging sessions. These graphs show the difference in performance between the training and the testing sets for each pair of imaging sessions. **(b)** The relationship between the difference in performance from **a** and the number of sessions between each pair in the forward direction (upper triangle outlined in **a**). Black lines show best linear fit. **(c)** The difference in Z_diff_ between neurons tracked between pairs of imaging sessions for conditioned and *pseudo*-conditioned mice. **(d)** The relationship between difference in Z_diff_ and number of sessions between each pair in the forward direction (upper triangle outlined in **c**). **(e)** Similarity of the Z_diff_ scores of neurons tracked between pairs of sessions were assessed using Pearson correlation (*ρ*). The panels show the similarity between each pair of sessions averaged across conditioned (left) and *pseudo*-conditioned mice (right). Black lines show best linear fit. Statistics: Spearman’s rank correlation, ^†^p < 0.1, *p < .05, **p < .01, ***p < 0.001, ^n.s.^p > .05.

**Figure S7:**
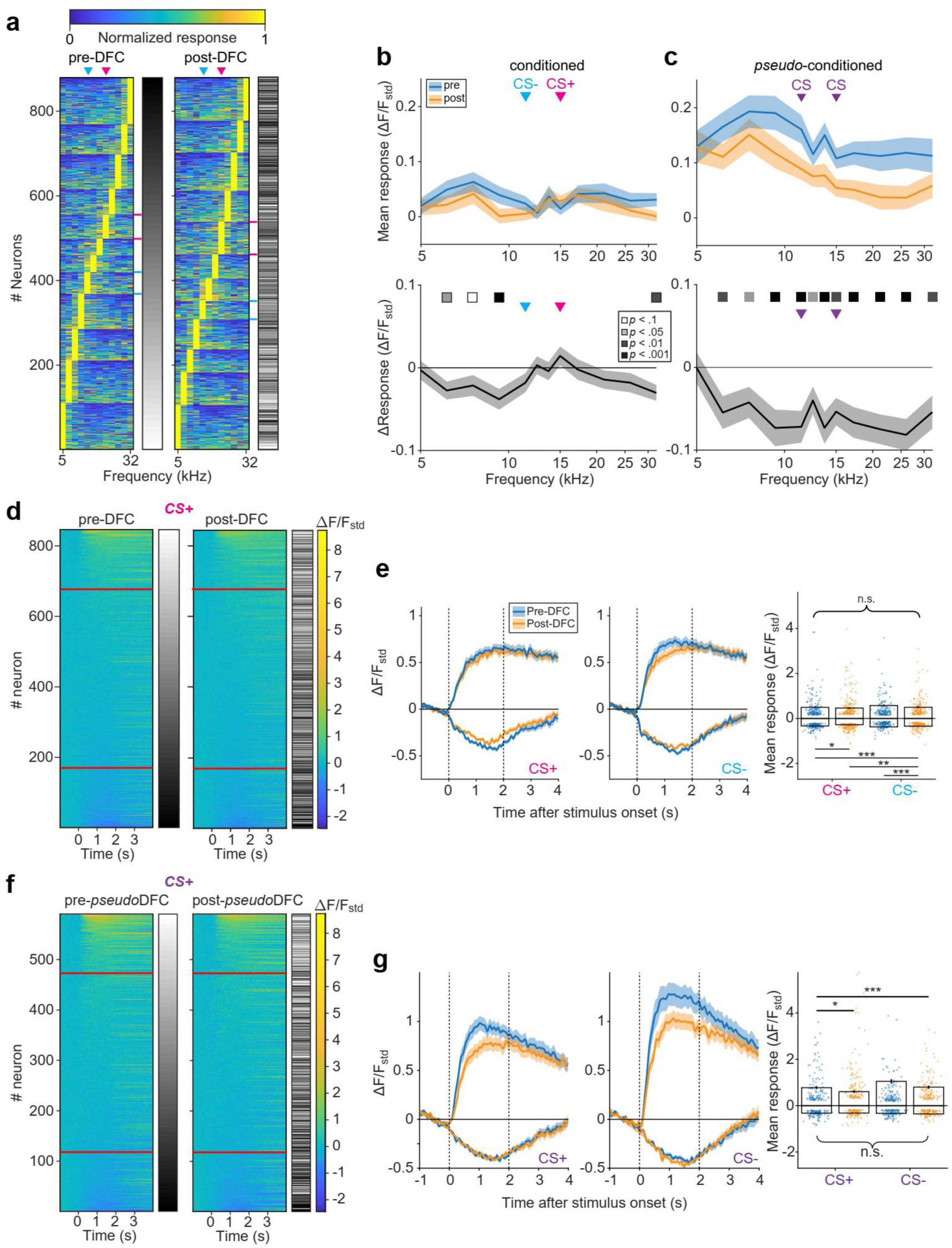
Changes in frequency response after conditioning. **(a)** Normalized frequency response curves of all neurons responsive at least once pre-DFC (imaging sessions 1-4) and once post-DFC (imaging sessions 5-8). Responses from neurons present in more than one session pre- or post-DFC are averaged together. Neurons are ordered according to their best frequency and secondarily by sparseness. The grey bars indicate the identity of each neuron pre- and post-DFC. The bars indicate the neurons with CS- (cyan) and CS+ (magenta) best frequencies. **(b)** (Upper panel) The mean frequency response curve of all responsive neurons (*not* normalized) pre- and post-DFC. (Lower panel) Change in frequency response curve across responsive all neurons. **(c)** Same as **b** for *pseudo*-conditioned mice. Statistics: paired *t*-test, *p*-values corrected for multiple comparisons by false-discovery rate. **(d)** Mean CS+-responses ordered by magnitude from conditioned mice (*N* = 14 mice) pre-DFC (left) and post-DFC (right). The grey bars indicate the identity of each neuron pre- and post-DFC. Red lines indicate the upper and lower 20^th^ percentiles. **(e)** Mean responses to CS+ and CS- of the upper and lower quartile of neurons ordered by response magnitude in **d**. **(f)** Same as **d** but for responses to CS+ from *pseudo*-conditioned mice. **(g)** Same as **e** but for responses to the CS+ and CS- in *pseudo*-conditioned mice. 2-way ANOVA with Tukey’s multiple comparisons test. Data are shown as mean ± SEM. See Tables S14, S15, S17 & S18 for full statistical results. ^†^p < 0.1, *p < 0.05, **p < 0.01, ***p < 0.001, ^n.s.^p > 0.10.

**Figure S8:**
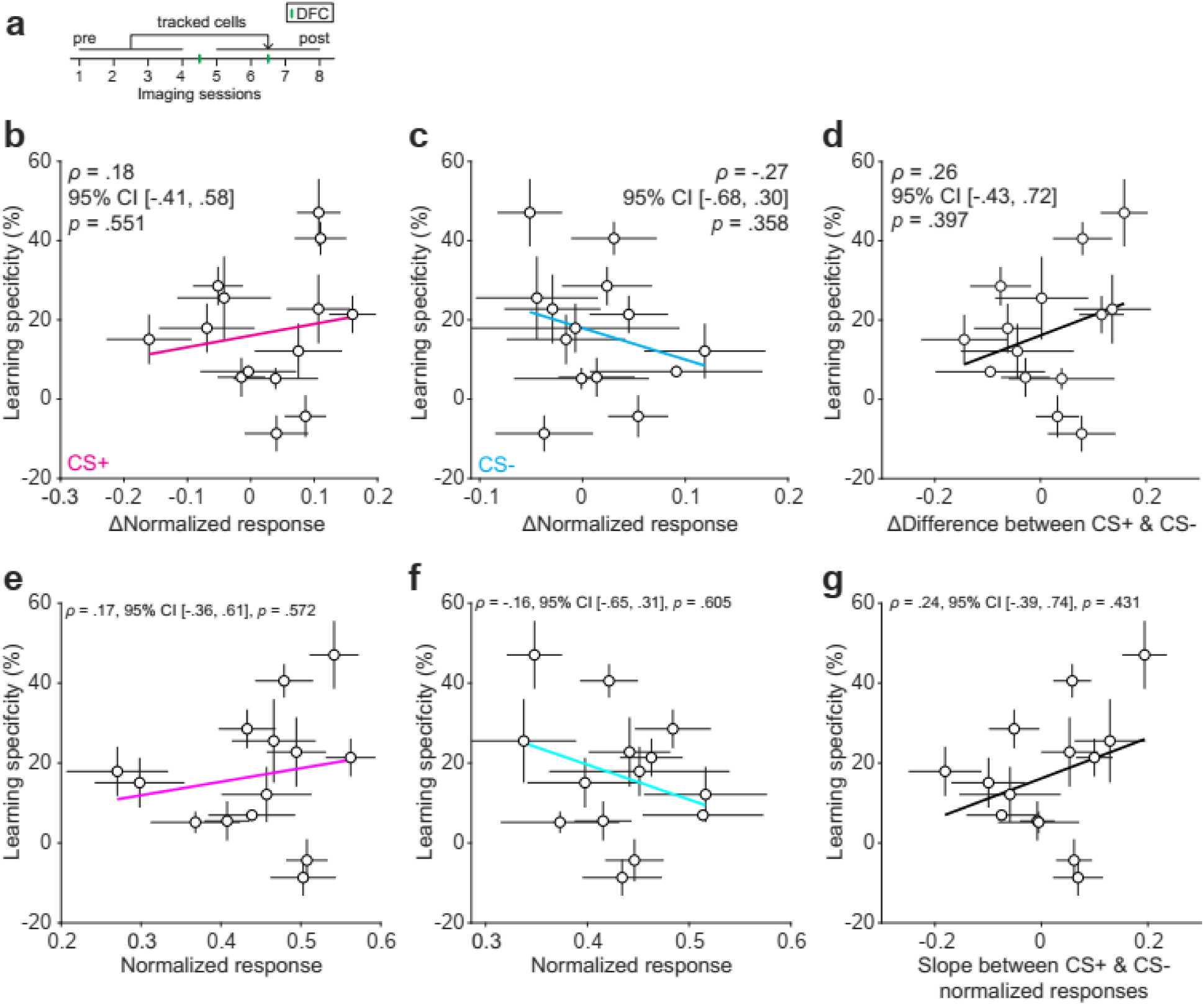
Changes in normalized responses and normalized responses post-DFC do not correlate with learning specificity. **(a)** Cells included were responsive in at least one imaging session pre- (sessions 1-4) and post-DFC (sessions 5-8). **(b)** Change in normalized response magnitude to CS+ post-DFC against mean learning specificity across retrieval sessions 1-4. Magenta line represents the best linear fit. **(c)** Same as **b** but for change in response to CS-. **(d)** Same as **b** but for change in difference between CS+ and CS-. **(e)** Normalized response magnitude to CS+ post-DFC against mean learning specificity across retrieval sessions 1-4. Magenta line represents the best linear fit. **(f)** Same as **e** but for response to CS-. **(g)** Same as **e** but for difference in normalized response magnitude between CS+ and CS-. Statistics: Spearman’s correlation (*N* = 14). Data are shown as mean ± sem.

**Figure S9:**
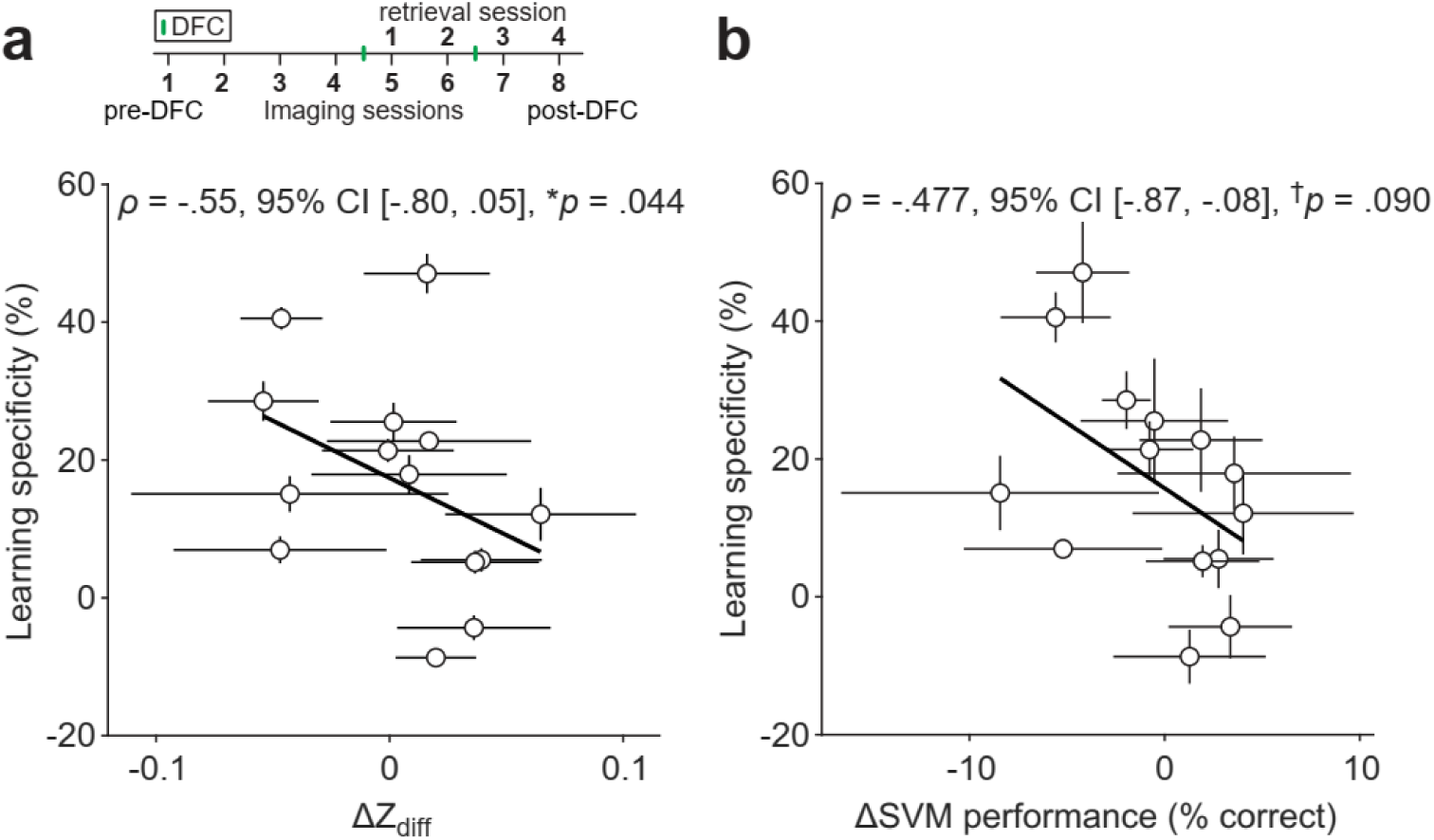
Changes in neuronal discriminability negatively correlate with learning specificity. **(a)** Change in neuronal discriminability was calculated as the difference between the mean discriminability across pre-DFC imaging sessions (1-4) and the mean discriminability across post-DFC imaging sessions (5-8). Change in Z_diff_ from pre- to post-DFC against mean learning specificity post-DFC (retrieval sessions 1-4). Black line represents the best linear fit. **(b)** Same as **a** but for change in SVM performance. Data are shown as mean ± sem. Statistics: Spearman’s correlation. ^†^*p* < 0.1, \**p* < .05, \*\**p* < .01, \*\*\**p* < 0.001, ^n.s.^*p* > 0.10.

**Figure S10:**
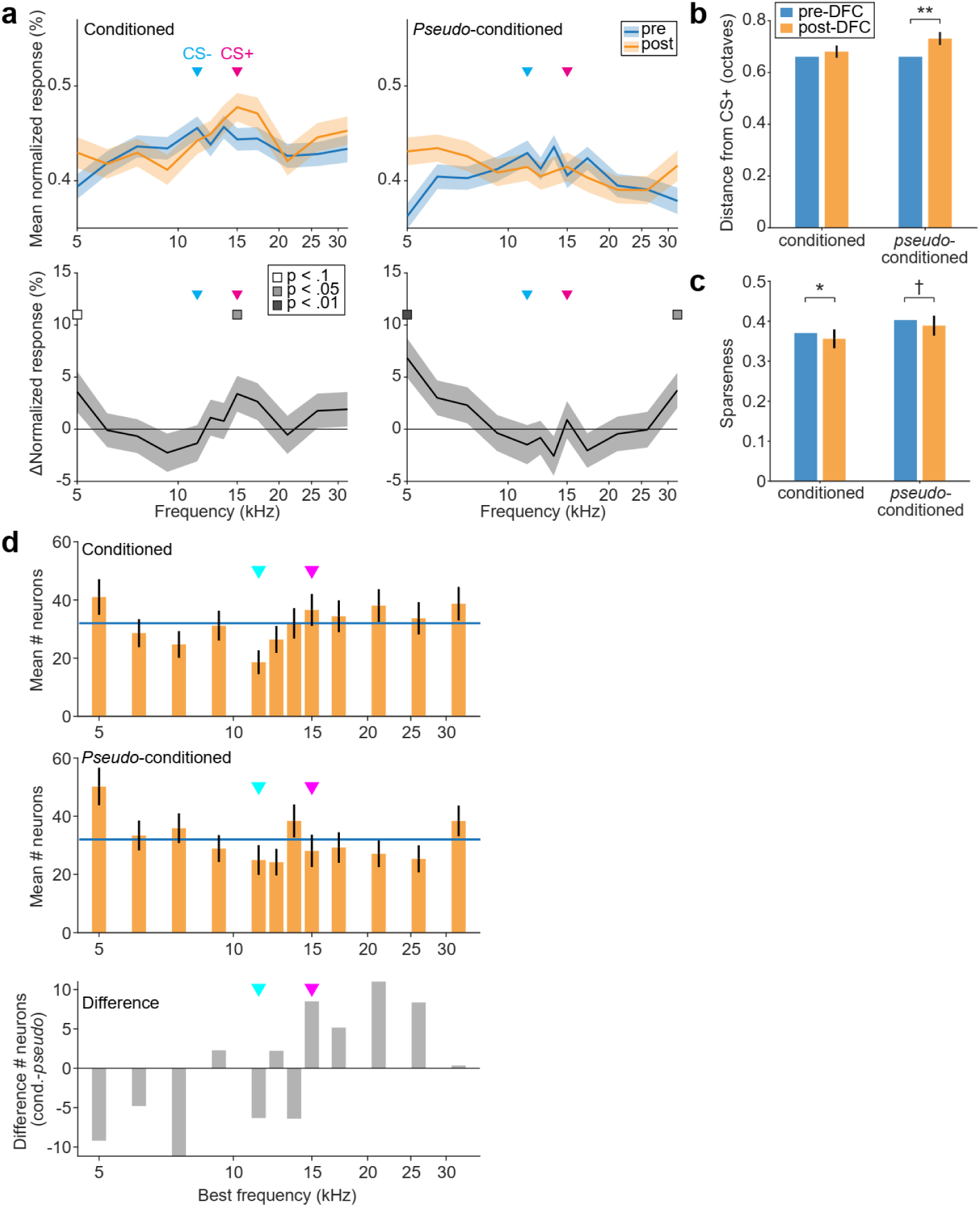
Effects of best frequency distributions on changes in response post-DFC. **(a)** The minimum number of neurons with best frequency at each tested frequency pre-DFC (n = 32) across conditioned and *pseudo*-conditioned mice was resampled with replacement (x250) from populations of neurons with best frequency at each frequency in conditioned and *pseudo*-conditioned mice. This had the effect to normalize the pre-DFC frequency distributions across neurons from the conditioned and *pseudo*-conditioned mice. The top panels show the mean normalized responses pre- (blue) and post-DFC (orange) for conditioned and *pseudo*-conditioned mice. The bottom panels show the % change in normalized response for the two groups. **(b)** Mean distance of the best frequency from CS+ pre- and post-DFC, using the same resampling as in **a**. **(c)** Sparseness of frequency tuning pre- and post-DFC, using the same resampling as in **a**. **(d)** Mean best frequency distributions post-DFC of the resampled neurons from **a** for conditioned (top) and *pseudo*-conditioned (middle) mice. The pre-DFC distribution is indicated by the blue line. The bottom panel shows the difference between the post-DFC best frequency distributions of the conditioned and *pseudo*-conditioned mice. Significance p-values indicate the percentile of the shuffled distributions at which zero occurred for the difference between the pre- and post-DFC for each frequency (**a**), distance of best frequency from CS+ (**b**) and sparseness (**c**). Error bars in all panels: ± sd of resampled data.

**Figure S11:**
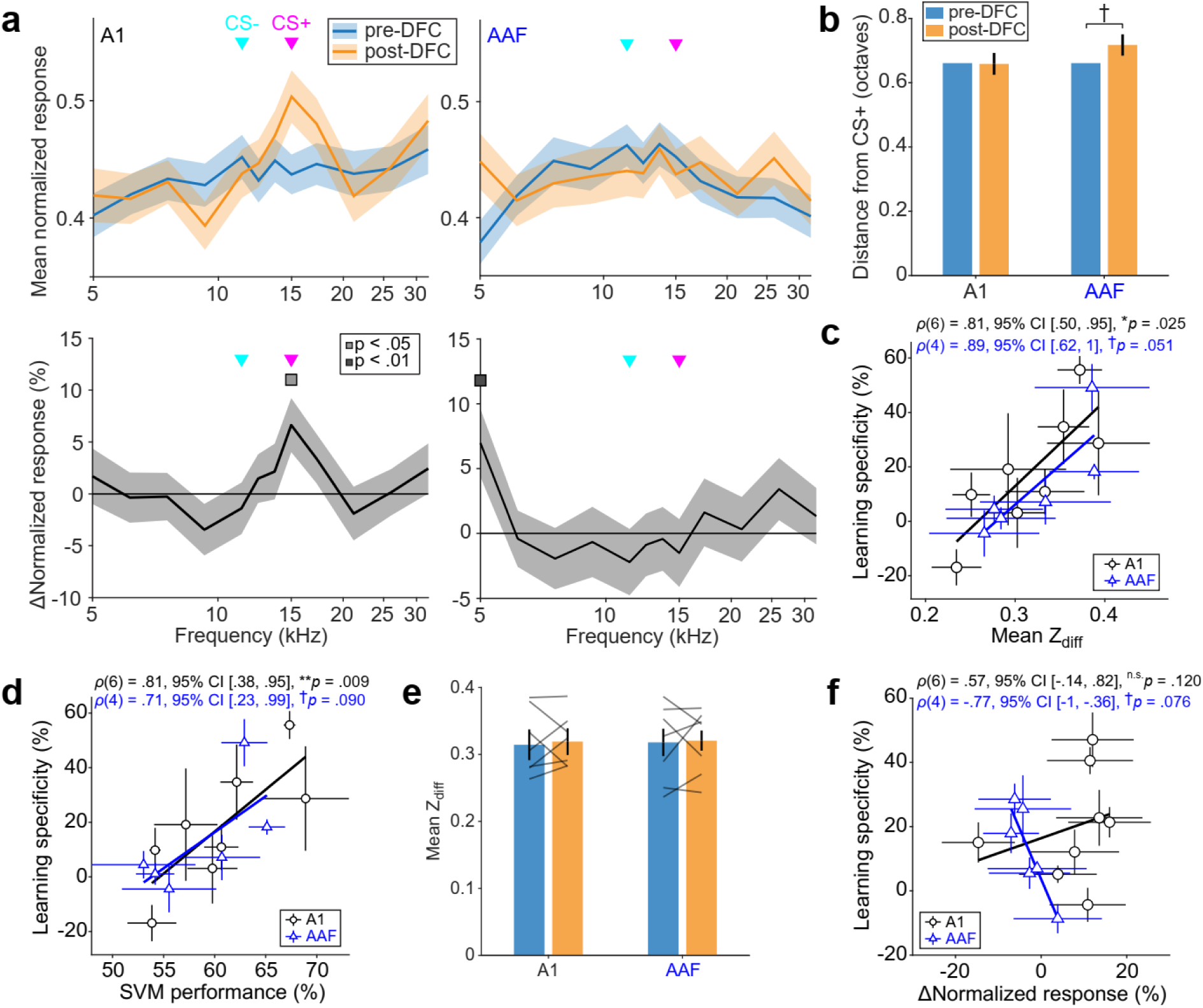
Effects of region of sampling on changes in response post-DFC and prediction of learning specificity. **(a)** The minimum number of neurons with best frequency at each tested frequency pre-DFC (n = 16) across conditioned mice with imaging regions in putative A1 or AAF was resampled with replacement (x250) from populations of neurons with best frequency at each frequency. This had the effect to normalize the pre-DFC frequency distributions across neurons from imaging windows in A1 or AAF. The top panels show the mean normalized responses pre- (blue) and post-DFC (orange) for A1 and AAF in conditioned mice. The bottom panels show the % change in normalized response for the two groups. **(b)** Mean distance of the best frequency from CS+ pre- and post-DFC, using the same resampling as in **a**. **(c)** Relationship between mean Z_diff_ (± sem) across pre-DFC imaging sessions (1-4) and learning specificity (± sem) in retrieval session 1 for conditioned mice with imaging regions in putative A1 (black) and AAF (blue). **(d)** Relationship between mean SVM performance across pre-DFC imaging sessions (1-4) and learning specificity (± sem) in retrieval session 1 for conditioned mice with imaging regions in putative A1 (black) and AAF (blue). **(e)** Change in mean Z_diff_ (± sem) from pre- to post-DFC for conditioned mice with imaging regions in A1 and AAF. **(f)** Relationship between change in normalized response at CS+ (± sd) and mean learning specificity (± sem) post-DFC (retrieval sessions 1-4). Significance p-values in **a** and **b** indicate the percentile of the shuffled distributions at which zero occurred for the difference between the pre- and post-DFC for each frequency (**a**) and distance of best frequency from CS+. Error bars in **a** and **b**: ± sd of resampled data from **a**.

**Figure S12:**
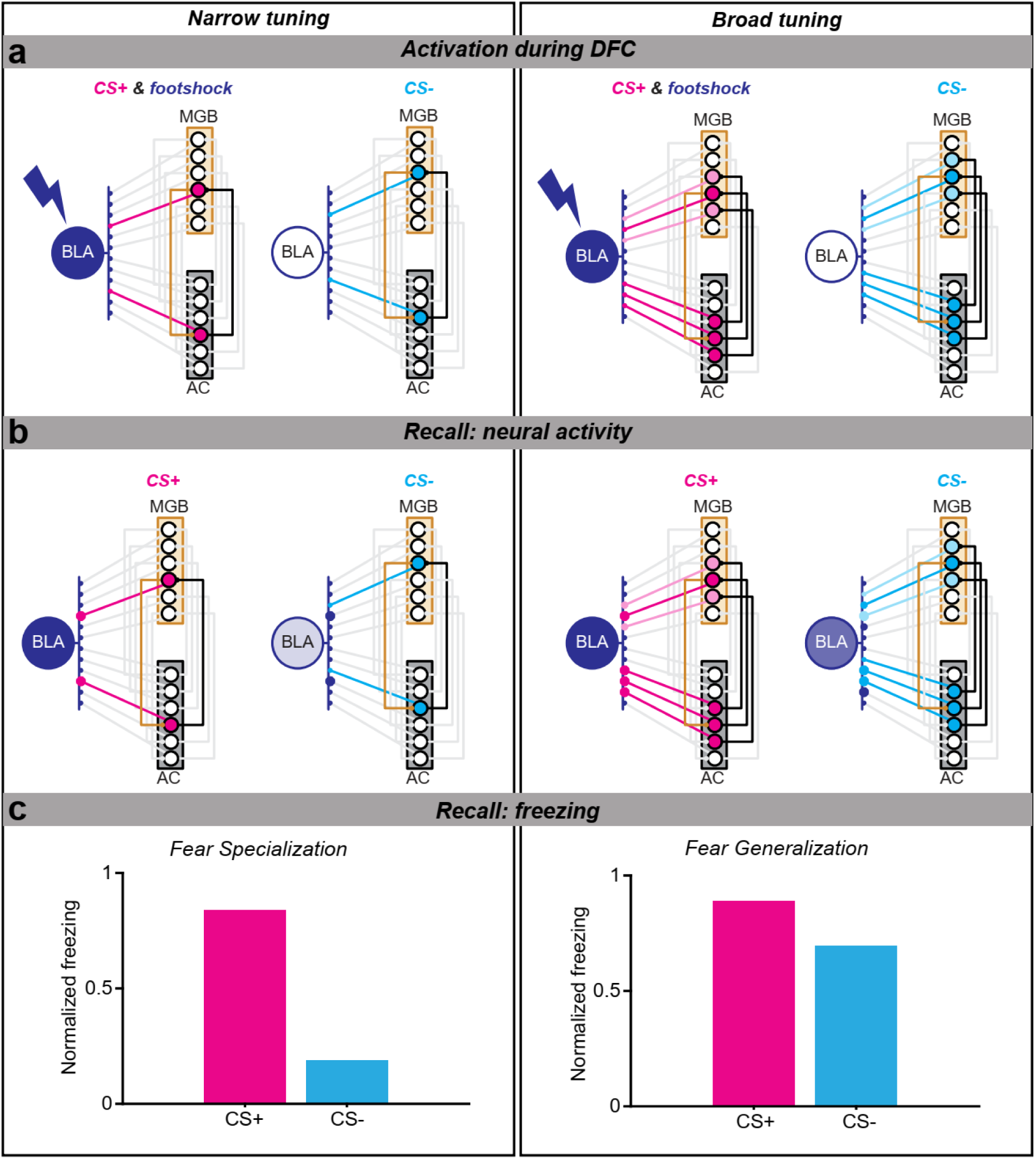
Model schematic. The stages of the model are outlined with narrow tuning in AC (high discriminability, left column) and broad tuning (low discriminability, right column). **(a)** Activation of projections during DFC. MGB is narrowly tuned and provides input to AC (orange connections). Feedback from AC to MGB (black connections) is narrow or broad depending upon the tuning width of AC, e.g. in different subjects. A foot-shock (lightning shape) is delivered simultaneously with the CS+ (magenta) and activates the BLA. CS- (cyan) is presented alone and there is no activation of BLA. The weights of the connections between AC and BLA, and MGB and BLA are strengthened depending on their co-activation of BLA. Feedback connections from AC help to strengthen the MGB projections to BLA. Connections with increased weights are shown as larger circles. **(b)** During memory recall, with narrow AC tuning, CS+ stimulus activates consolidated connections, thus activating BLA, whereas the CS- does not. With broad AC tuning, both CS+ and CS- activate consolidated connections, leading to BLA activation with both stimuli. **(c)** The normalized freezing output of the model (relative levels of BLA activation). Broad tuning leads to increased levels of fear generalization.

**Figure S13:**
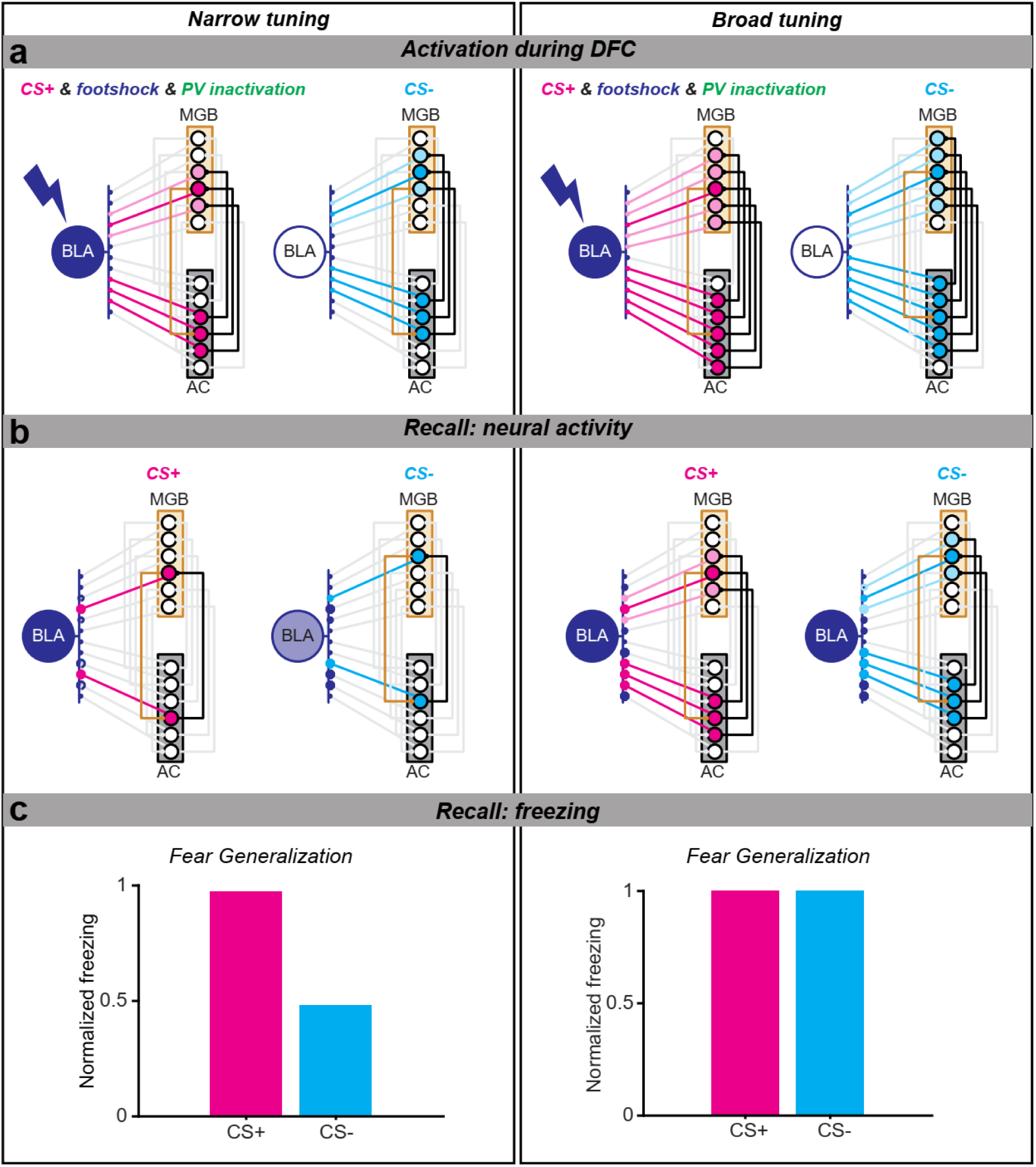
Model schematic with PV inactivation of AC during DFC. The process of the model is the same as Fig. S9. PV inactivation is modeled by lowering cortical inhibition. It effectively increases the tuning width of AC tuning (therefore lower discrimination in AC). Following activation (**a**), AC tuning is returned to its original narrow and broad tuning (as in Fig S11). Even with narrow tuning, CS- now activates strengthened AC-BLA and MGB-BLA connections (**b,** left panel) leading to increased fear generalization (**C,** left panel) compared with no PV inactivation (Fig S11). With broad tuning, both CS+ and CS- activate many strengthened AC-BLA and MGB-BLA connections (**b**, right panel) leading to high levels of fear generalization compared with no PV inactivation (**c**, right panel).

**Figure S14:**
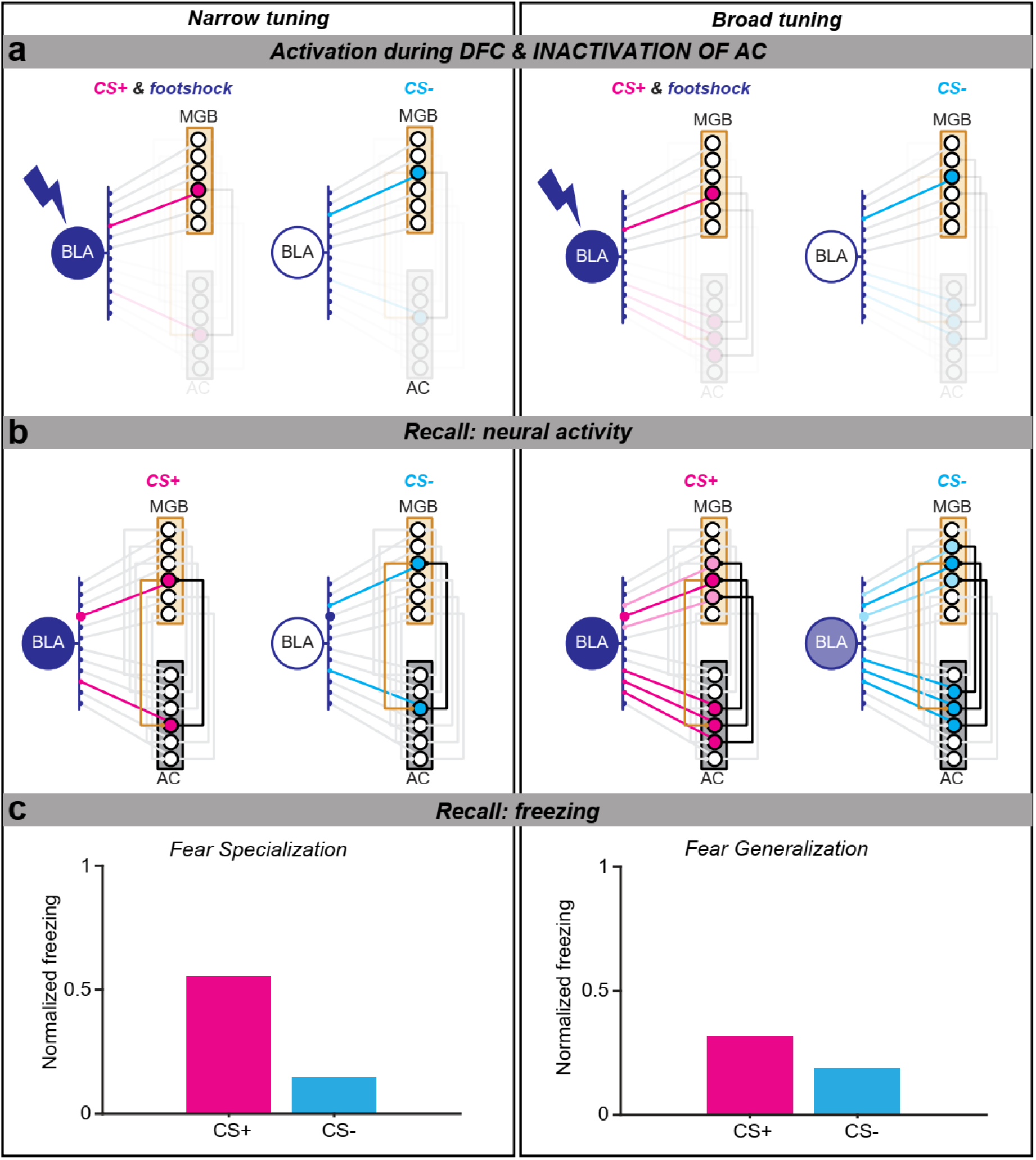
Model schematic with AC inactivation during DFC. The process of the model is the same as Fig. S9. AC inactivation is modeled by setting cortical currents to zero during the conditioning phase. Following inactivation (**a**), AC tuning is returned to its original narrow and broad tuning (as in Fig. S11). **(b)** Schematic of recall with AC active. **(c)** Freezing to CS+ and CS- presentations during recall. Although freezing is lower in general, there is still discrimination in both cases but much lower than when AC is active throughout.

**Figure S15:**
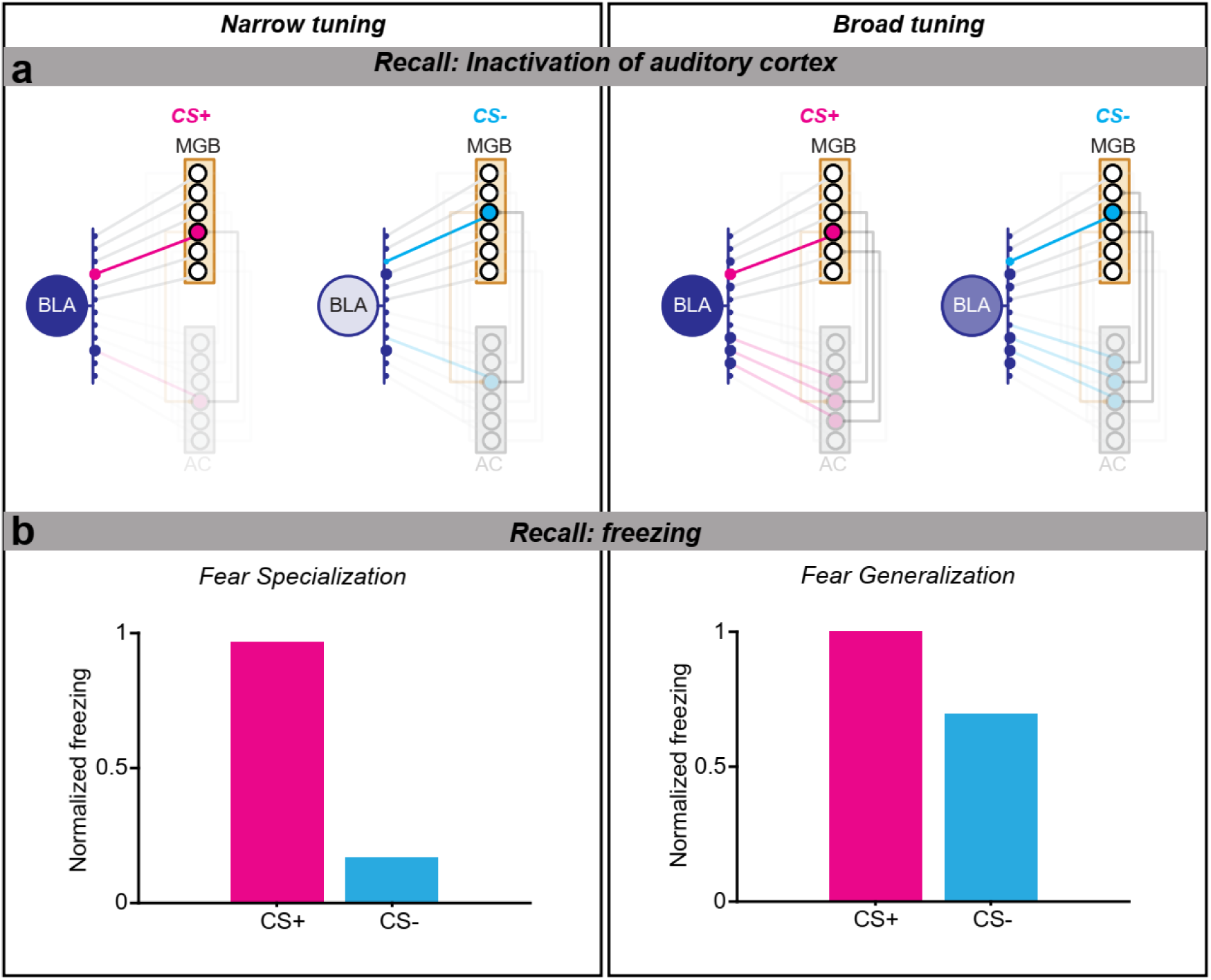
Model schematic with AC inactivation during memory recall. Activation and consolidation during DFC are the same as Fig S11. **(a)** shows activation of MBG and BLA without the presence of AC. With broad tuning, CS- still activated strengthened MGB-BLA connections thus increasing BLA activation. **(b)** Results of freezing during memory recall are very similar to when there are no interventions (Fig. S11).

**Figure S16:**
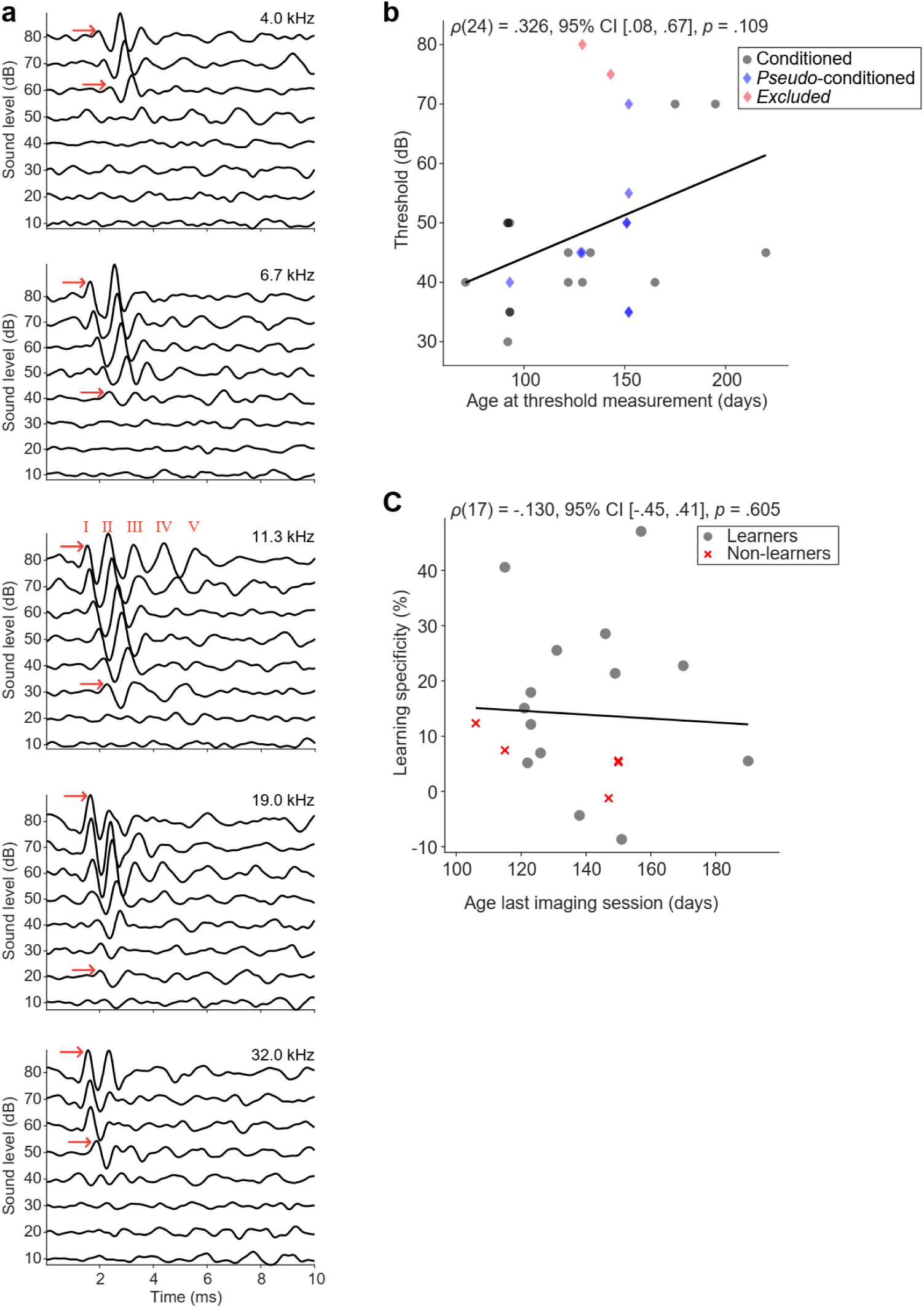
Auditory Brainstem Responses. **(a)** Example ABR responses to 5 frequencies presented at 7 different levels. **(b)** Relationship between age at ABR threshold measurement and threshold (mean of frequencies closest to CS+ and CS-). Mice with threshold greater than 70dB (red) were excluded from the study. **(c)** Relationship between age at the last imaging session and learning specificity. Black lines in **b** and **c** are best linear fits. Statistics: Spearman’s rank correlation.

Table S1: Statistics for all figures.

Table S2: Statistics comparing normalized responses at CS+, CS- and CSc.

Table S3: Statistics comparing absolute normalized responses at CS+, CS- and CSc.

